# Generation of human appetite-regulating neurons and tanycytes from stem cells

**DOI:** 10.1101/2024.07.11.603039

**Authors:** Zehra Abay-Nørgaard, Anika K Müller, Erno Hänninen, Dylan Rausch, Louise Piilgaard, Jens Bager Christensen, Sofie Peeters, Alrik L. Schörling, Alison Salvador, Viktoriia Nikulina, Yuan Li, Janko Kajtez, Tune H Pers, Agnete Kirkeby

**Author notes:** These authors contributed equally Correspondence.

## Abstract

The balance between energy intake and expenditure is controlled by the hypothalamus, a small brain region characterised by high neuronal diversity. Specifically, the arcuate nucleus (ARC) and ventromedial hypothalamus (VMH) are key hypothalamic nuclei controlling appetite through behavioural response to circulating humoral signals. Yet, despite their physiological importance, the cellular and functional characteristics of this highly specialised neural region has been studied mainly in animals due to a lack of human models. Here, we fine-tuned the differentiation of human pluripotent stem cells toward the ARC and VMH hypothalamic nuclei and identified key subtype-specific progenitor markers of these subregions. We demonstrate that the timing for initiation and termination of bone morphogenetic protein (BMP) signalling is essential for controlling subregional specification of tuberal hypothalamic progenitors along the anterior-posterior axis, balancing VMH versus ARC fates. A particular population of SHH^-^/NKX2.1^+^/FGF10^high^/RAX^high^/TBX3^high^ posterior tuberal progenitors was identified as the source for generation of ARC-associated agouti-related peptide (AGRP) neurons and tanycytes whilst anterior tuberal SHH^+^/NKX2.1^+^/FGF10^low^/RAX^low^/TBX3^low^ progenitors generated VMH phenotypes including NR5A1 neurons. Upon maturation *in vitro* and in xenografts, ARC-patterned progenitors gave rise to key appetite-regulating cell types including those producing AGRP, prepronociceptin (PNOC), growth hormone-releasing hormone (GHRH), thyrotropin-releasing hormone (TRH) and pro-opiomelanocortin (POMC), as well as tanycyte glial cells. Differentiated ARC cultures showed high transcriptomic similarity to the human ARC and displayed evidence of functionality by AGRP secretion and responsiveness to leptin and fibroblast growth factor 1 (FGF1). In summary, our work provides insights into the developmental lineages underlying hypothalamic subregional specification and enables access to highly characterised human ARC and VMH cultures, which will provide novel opportunities for investigating the cellular and molecular pathways triggered by obesity-associated genetic variants and weight-regulating stimuli.

## Introduction

Food intake and blood glucose homeostasis is heavily controlled by hypothalamic circuits, especially by so-called first-order ARC neurons, AGRP and POMC, which integrate and relay information about an organism’s fed state to second-order neurons deeper into the hypothalamus and other brain regions^1–3^. The neighbouring VMH plays a similarly important role as the satiety centre of the brain by sensing glucose levels and suppressing food intake^4,5^. In addition, non-neuronal cells, in particular tanycytes, which are specialised radial glial-like cells lining the third ventricle of the ARC, play a key role in the relay of humoral cues into the hypothalamus^6–8^. Consequently, dysregulation of these hypothalamic cell populations is coupled to metabolic diseases such as obesity and type 2 diabetes^2^. In line with this, appetite-regulating hormones such as leptin^9^ as well as recent appetite-lowering drugs for obesity targeting the glucagon-like peptide 1 receptor (GLP1R), including semaglutide and tirzepatide, are hypothesised to act on hypothalamic circuits, the mechanism of which is only partially resolved^10–13^.

While the development, cellular composition and molecular function of the hypothalamus has been studied in the mouse and other animal models^14,15^, the means to study the human hypothalamus have been limited due to its inaccessibility. Consequently, functional *in vitro*-derived hypothalamic neurons from human pluripotent stem cells constitute a unique tool to study the molecular pathways involved in appetite regulation, obesity, and type 2 diabetes^16,17^ as well as for targeted drug discovery^18,19^. However, generating authentic cultures of subregional hypothalamic neurons from stem cells remains a challenge due to the high complexity of the hypothalamic nuclei. Furthermore, the lineage relationship between each neural progenitor and its corresponding adult neuronal subtypes is unclear as the regional anatomical partitioning during development and the occurrence of potential inter-regional neuronal migration remains highly debated^20–24^. Attempts to recreate complex hypothalamic developmental patterning *in vitro* with human pluripotent stem cells (hPSCs) have generally resulted in cultures of high neuronal heterogeneity containing cell types from various different hypothalamic and non-hypothalamic regions, and a low yield or absence of important cells types belonging to the ARC and VMH, including the key appetite-regulating AGRP, prepronociceptin (PNOC), growth hormone-releasing hormone (GHRH) and NR5A1 neurons as well as tanycytes^17,25,26^. Access to reproducible and well-controlled human subregional hypothalamic *in vitro* models would thereby address a critical unmet need in metabolic research and allow for high-throughput studies on the cellular and molecular effects of genetic variants, metabolic compounds, hormones and drug candidates relevant for obesity and metabolic disease.

Here, we developed novel differentiation protocols for human pluripotent stem cell (hPSCs) and showed that we could generate POMC, AGRP, GHRH, PNOC, NR5A1 and tanycyte lineages by recapitulating early tuberal hypothalamic patterning events identified in the chick^27^. We found that timed addition of BMP was crucial for controlling patterning towards posterior tuberal progenitors giving rise to ARC-related AGRP neurons and tanycytes versus anterior tuberal progenitors giving rise to VMH-related NR5A1 neurons. We further validated the functionality of our ARC cultures through calcium imaging, AGRP secretion and leptin responsiveness and we confirmed authentic lineage commitment of the progenitors through *in vivo* transplantation to rats. In summary, these novel human *in vitro* models of the ARC and VMH nuclei provide important new insights into subregional hypothalamic development and enable a novel platform for the molecular dissection of appetite control, including possibilities for drug screening and large-scale genetic perturbation studies.

## Results

### *In vitro* model of human ARC

To produce human neurons from specific subregions of the hypothalamus, we subjected human embryonic stem cells (hESCs) to a multitude of neural differentiation paradigms, testing various timings and concentrations of molecules targeting the key neurodevelopmental pathways WNT, BMP and sonic hedgehog (SHH) (**Fig. 1a**). Each differentiation (n = 113) was analysed by quantitative reverse transcriptase PCR (qRT-PCR) using a primer panel of 64 genes marking the hypothalamus and its surrounding regions. To identify combinations of factors that produced distinct hypothalamic fates, we applied Principal Component Analysis (PCA), grouping the differentiation outcomes based on their gene expression profiles (**Supplementary** Fig. 1a). Interestingly, PC1 separated samples based on the expression of canonical ARC and paraventricular nucleus (PVN) genes. We next investigated how different morphogens, added at various time points during differentiation, could drive regionalisation towards ARC versus PVN fates (i.e., along the PC1 axis). Covariant analysis revealed that early addition of CHIR99021 (an activator of canonical WNT signalling) at day 0 (d0) and late addition of SHH at d6 favoured the derivation of PVN fates (**Supplementary** Fig. 1b). Based on this information, we tested the validity of the analysis by designing a predicted optimal PVN protocol (**Supplementary** Fig. 1c) and confirmed the generation of cultures containing PVN-specific OTP/*SIM1* and *BRN2* progenitors, which upon maturation gave rise to thyrotropin releasing hormone (TRH) and Corticotropin releasing hormone (CRH) neurons (**Supplementary** Fig. 1d-f). In contrast, later addition of CHIR at d2 was predicted to favour ARC specification, but only when combined with the addition of BMP (**Supplementary** Fig. 1b). Indeed, replicate experiments and *in situ* hybridisation (ISH) confirmed that the addition of BMP in early stages of the protocol was essential for induction of key ARC transcription factors *RAX* and *TBX3* (**Fig. 1b and Supplementary** Fig. 1e) as well as for the later generation of mature neurons expressing α-melanocyte stimulating hormone (α-MSH, derived from the POMC genes) and AGRP (**Fig. 1b**) – secreted neuropeptides which are centrally involved in controlling food intake in mammals^1,2^. From this information, we designed a predicted optimal ARC differentiation protocol involving timed activation of SHH, WNT and BMP pathways (**Fig. 1c**). We also added IGFBP3 - a developmentally secreted protein we identified to be expressed in the pre- and post-mitotic populations of *TBX3*+ ARC precursors in a human fetal hypothalamic dataset (**Supplementary** Fig. 1g-i) ^28^, and which appeared beneficial for the generation of POMC and AGRP neurons (**Supplementary** Fig. 1j).

**Fig. 1.**
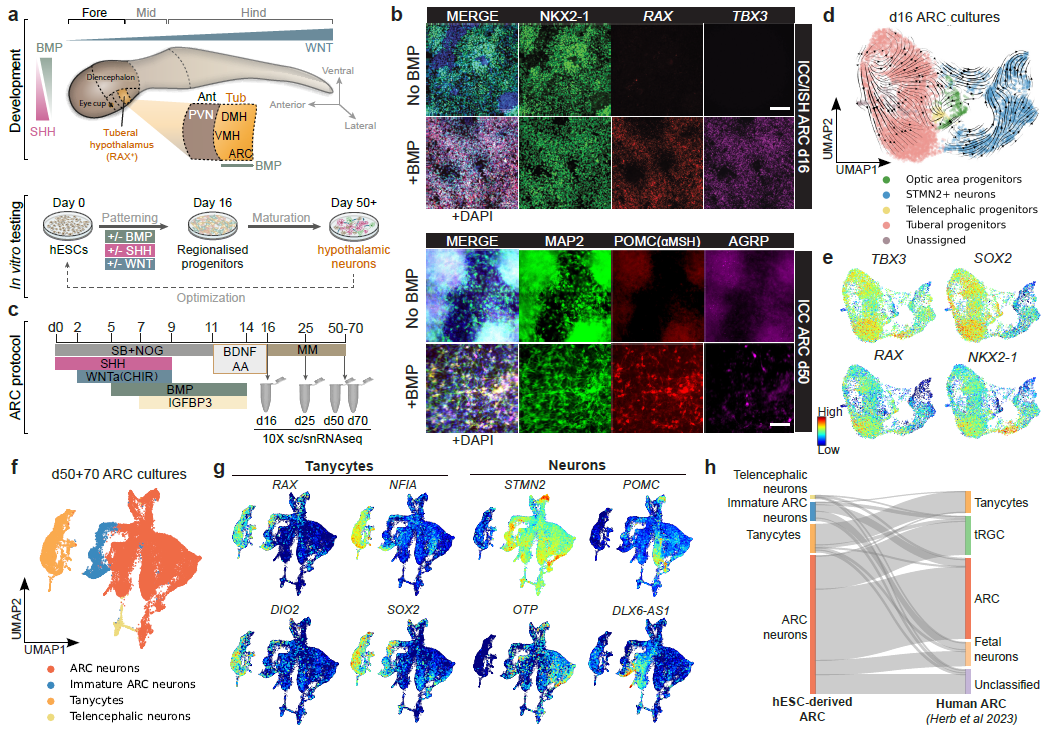
hESC differentiation protocol generates NKX2-1+/*RAX*+/*TBX3*+ progenitors that give rise to ARC-specific neurons and tanycytes. **a**, Top: Schematic representation of the developing neural tube and local growth factor gradients involved in hypothalamic patterning. Middle: Schematic representation of test conditions for *in vitro* hESC differentiation by manipulating morphogens and growth factors. **b**, Top: Combinatorial ICC (NKX2-1) and *in situ* hybridization (ISH) (*RAX* and *TBX3*) (ICC/ISH) on d16 ARC progenitors differentiated with and without BMP addition. Bottom: ICC of MAP2, POMC (αMSH) and AGRP on d50 ARC neurons differentiated with and without BMP treatment. Scale bars, 100 μM. **c**, Schematic overview of the optimised ARC differentiation protocol. Samples were taken at day (d) 16, 25, 50 and 70 for sc/snRNAseq. **d**, UMAP of scRNAseq data from d16 ARC progenitors showing annotated clusters and RNA velocity. **e**, Feature plots of highly expressed early ARC genes in the d16 scRNAseq dataset. **f**, UMAP of snRNAseq data from the ARC protocol at d50 and d70 with annotated clusters. **g**, Feature plots of marker genes for tanycytes (*RAX, NFIA, DIO2, SOX2*) neurons (*STMN2*) and ARC-specific neurons (*OTP, POMC, DLX6-AS1*) in the d50 and d70 sn-RNAseq dataset. **h**, Sankey diagram showing mapping of hESC-derived cell types from the d50+70 dataset against a compiled annotated dataset of human fetal and adult hypothalamic cells^28^. AA, ascorbic acid; ARC, arcuate nucleus; BDNF, bone-derived neurotrophic factor; BMP, Bone morphogenic protein; DMH, dorsomedial hypothalamus; Fore, Forebrain; Hind, hindbrain; MM, maturation medium; Mid, midbrain; NOG, Noggin; PVN, paraventricular nucleus; SB, SB-431542; SHH, Sonic hedgehog; tRGC, transitional radial glial cells; VMH, ventromedial hypothalamus.

We next explored the cellular composition of our optimised ARC protocol cultures through single cell and single nucleus RNA sequencing (sc/snRNAseq) of three biological replicates at days 16, 25, 50 and 70 (**Supplementary** Fig. 2a). Analysis of the early-stage cultures at d16 showed high reproducibility between replicate batches with cultures mainly composed of *NKX2-1^+^/RAX^+^*/*TBX3*^+^ tuberal progenitors (70±1.4%, mean±SD) and postmitotic neurons (23±3.7%) with minor contaminating populations of *CRABP1*^+^/*VSX2*^+^/*NR2F1*^+^ optic area progenitors (3.4±2.9%), *FOXG1*^+^ telencephalic progenitors (1.3±0.3%), and some unassigned cells (1.9±0.2%) (**Figure 1d,e, Supplementary** Fig. 1k-m **and Extended Data Table 1**). Further maturation of the cells for 50 to 70 days under either 2D monolayer or 3D spheroid conditions produced cultures of ARC neurons with a minor cluster of contaminating telencephalic neurons and a larger cluster of non-neuronal cells expressing *RAX*, *NFIA, DIO2, SOX2,* and *CRYM*, corresponding to tanycytes (**Fig. 1f,g and Supplementary** Fig. 2b-d). To investigate the transcriptional similarity of our *in vitro* derived cells to the human ARC, we mapped the d50+*70 in vitro* dataset to a compiled dataset of the entire human hypothalamus from fetal and adult stages^28^. This analysis showed that the hESC-derived ARC neurons mapped predominantly to “ARC neurons” in the human dataset, with a subset mapping to the human fetal “tRGC” (transitional radial glial cell) cluster representing early ARC progenitors expressing *TBX3*, *RAX* and *SOX2* (**Figure 1h and Supplementary** Fig. 2e-h). Importantly, the prospective tanycyte cluster mapped almost exclusively to “tanycytes” in the human dataset, despite several other non-neuronal cells (i.e. astrocytes, pericytes, ependymal cells, endothelial cells etc.) being present in the human dataset (**Supplementary** Fig. 2g,h). Less than 2.1% of the hESC-derived ARC cells mapped to other neighbouring hypothalamic nuclei and non-neuronal cell types from the human reference dataset (**Supplementary** Fig. 2h), thereby confirming correctly specified ARC fate of our hESC-derived cultures.

### Neuronal diversity of human ARC cultures

To assess the neuronal subtype composition of the hESC-derived ARC cultures, we extracted the ARC neuron cluster from the d50+70 dataset and performed integration and subclustering (**Fig. 2a and Supplementary** Fig. 3b-d). From this, we annotated 10 neuronal subtypes associated with the ARC, including POMC, PNOC, AGRP, GHRH and TRH neurons, and we confirmed the presence of several of these neuronal subtypes by immunocytochemistry (ICC) in 2D and 3D cultures at d50 (**Fig. 2b-d and Supplementary** Fig. 3a). As expected from mouse and human data, a subset of cells in the AGRP cluster also co-expressed the neuropeptide somatostatin (SST)^28,29^ (**Supplementary** Fig. 3e). We further identified two clusters of PNOC-expressing neurons, GHRH^+^/PNOC^+^ and PNOC^+^/TAC3^+^, representing additional ARC neuron subtypes which in the mouse have been shown to have important functions in controlling feeding behaviour^30^. In addition to PNOC, the mature GHRH neurons also co-expressed *PROX1*, *FOXP2* and the receptor for GLP1, *GLP1R* (**Supplementary Fig.3e**) as previously described in mouse^31^. In addition to clusters like DLX-AS1^+^/FOXP2^+^ and OTP^+^/UNC13C^+^, the data further revealed three subclusters of POMC neurons: POMC^+^/PRDM12^+^/LEPR^+^, POMC^+^/NR5A2^+^/TRH^+^ and POMC^+^/CRABP1^+^/TRH^+^ (**Fig. 2a,b**).

**Fig. 2.**
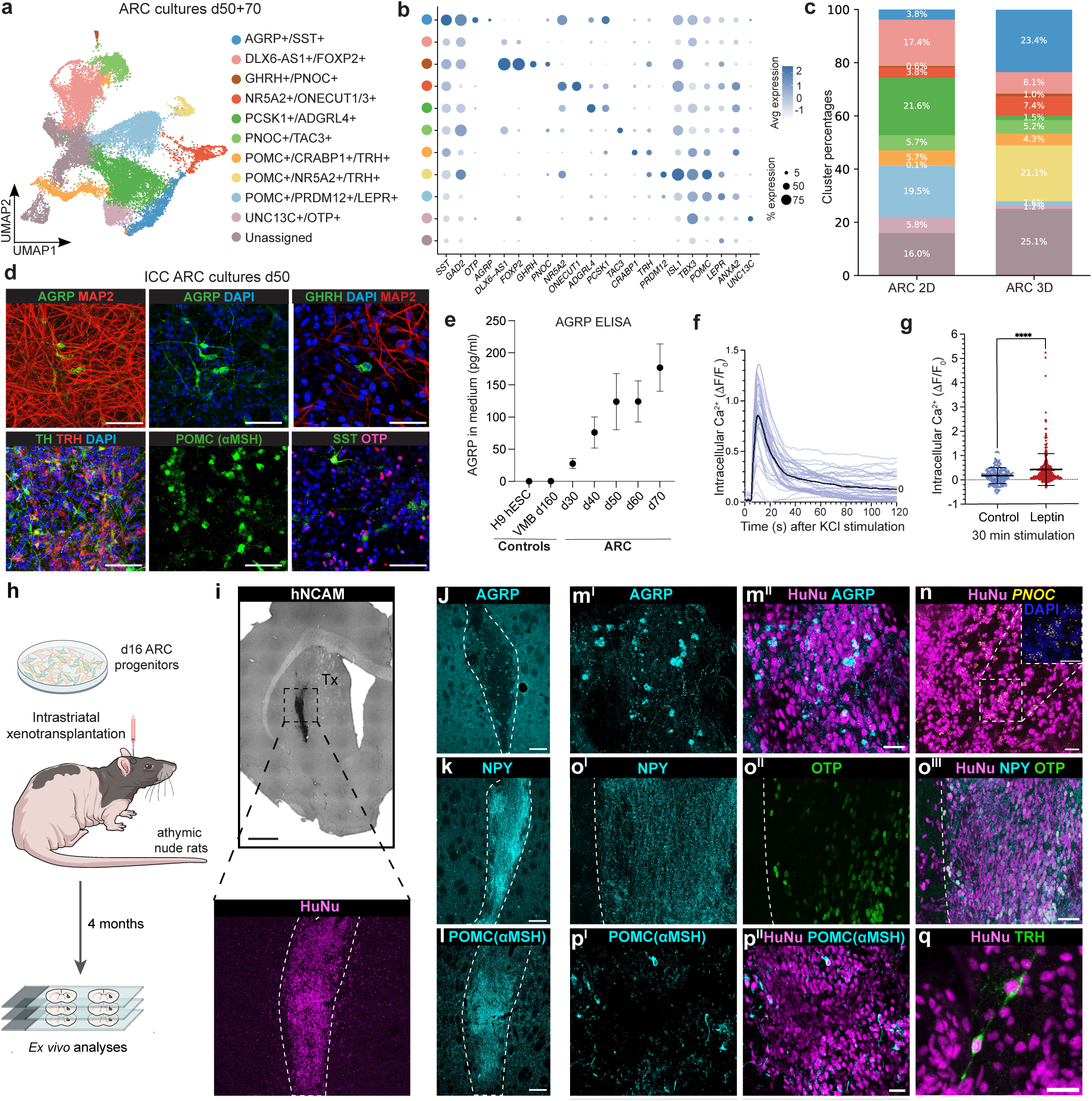
ARC progenitors mature to appetite-regulating neurons in 2D, 3D and in xenografts, and show responsiveness to leptin. **a**, UMAP of ‘ARC neurons’ cluster subsetted from day (d) 50 and d70 scRNAseq dataset (see Figure 1) with annotated clusters. **b**, Dot plot of key genes associated with ARC neuron clusters. **c**, Comparison of cluster percentages between 2D and 3D maturation. **d**, Immunocytochemistry (ICC) of 2D cultures at d50 depicting AGRP, GHRH, TRH, TH, POMC (αMSH), SST and OTP neurons. Scale bar, 50 µm. **e**, ELISA of secreted AGRP (pg/ml) detected in the culture medium (ARC: n=3 batches) compared to control (VMB, ventral midbrain organoid; hESC, human embryonic stem cells). **f**, Calcium imaging of d50 ARC cells stimulated with 50 mM KCl. Blue lines mark individual cells, black line marks average (n=35 cells). **g**, Calcium imaging of d50 ARC cultures after 30 min stimulation with 100 ng/mL leptin (control=169, leptin=304 neurons). Mann-Whitney, non-par-ametric test: ****p<0.0001. **h**, Schematic overview of ectopic xenotransplantation of d16 ARC progenitors into the striatum of athymic nude rats. **i**, Human graft of d16 ARC progenitors in the striatum of nude rats four months post-transplantation. Top: Human neural cell adhesion molecule (hNCAM) IHC staining. Scale bar, 1 mm. Bottom: human nuclear antigen (HuNu) immunofluorescence (IF) staining. Scale bar, 100 µm. **j-l**, IF images of AGRP (j), NPY (k) and POMC (αMSH) (l) within graft site (dotted lines). Scale bars, 100 µm. **m-q**, Close-up IF of AGRP (m^I+II^), PNOC (n) NPY (o^I^), OTP (o^II^), POMC (αMSH) (p^I+II^), and TRH (q) within the graft (with HuNu co-staining). Scale bars, 25 µm. ARC, arcuate nucleus.

The POMC/TRH co-expression was validated by ICC in 2D cultures (**Supplementary** Fig. 3f), and this subtype was also found to co-express *GLP1R* (**Supplementary Fig.3e**). Even though most neuronal subtypes were found in both 2D and 3D maturation conditions, the 3D maturation was particularly beneficial for generation of AGRP neurons, with this cluster comprising 23.4±1.18% (mean±SD) of all neurons in 3D compared to 3.8±0.79% in 2D (**Fig. 2c**). Another notable difference was a shift in proportions of POMC subtypes with the POMC+/NR5A2+/TRH+ subtype being enriched in 3D, possibly reflecting a shift to more mature phenotypes under spheroid conditions. We proceeded to perform functional assessment of the ARC neuronal cultures through various assays. Analysis of the culture medium by ELISA revealed AGRP peptide release from the cultures over time as the neurons progressively matured (**Fig. 2e**). Furthermore, electrophysiological stimulation of the neurons with potassium chloride (KCl) produced a robust response by calcium imaging, indicating that the neurons were capable of responding to an action potential stimulus (**Fig. 2f**). Lastly, we tested the ability of the cultures to respond to the leptin, a circulating hormone well-known to regulate appetite through action on leptin receptors (LEPR) in the hypothalamus^9^. Indeed, we confirmed the expression of *LEPR* in our ARC cultures (**Fig. 2b**), and we found that stimulation with leptin for 30 minutes caused a significant calcium response in the cultures, indicating functional leptin responsiveness (**Fig. 2g**).

To test the lineage fate commitment of the early d16 tuberal progenitors in our cultures, we performed intracerebral xenotransplantation of d16 NKX2.1^+^/RAX^+^/TBX3^+^ cultures to the adult striatum of immunodeficient athymic rats to allow the progenitors to mature in an ectopic *in vivo* environment without exogenously provided growth factors. The human xenografts were analysed at four months post-transplantation and characterised with respect to ARC neuron subtypes through a combination of immunofluorescence and ISH (**Fig. 2h,i**). The transplanted ARC progenitors survived and matured into AGRP, POMC, PNOC, OTP, TRH, and PNOC neurons (**Fig. 2j-q**). We also observed a strong NPY fibre staining in the graft closely resembling the diffuse NPY fibre innervation found in the endogenous rat ARC and PVN regions, which was distinctly different from the NPY+ interneuron cell bodies found in cortex and striatum (**Fig. 2k,o and Supplementary** Fig. 3i). This data confirmed that the hESC-derived tuberal progenitors were committed towards ARC neuron fate already at d16 of differentiation, and that the cells matured to a similar diversity of ARC neurons under *in vitro* and *in vivo* conditions. In summary, we show that our tuberal progenitor have the potential to develop into functional ARC-specific populations involved in energy homeostasis and appetite regulation, including neurons expressing AGRP, NPY, POMC, PNOC and GHRH.

### *In vitro* derived tanycytes respond to FGF1

In addition to neurons, our ARC cultures also contained a substantial population of tanycytes. To investigate the developmental trajectories of the tanycyte lineage, we integrated all cells from the d16 scRNAseq dataset with the tanycyte-annotated cluster from d25 and d50+70 (**Fig. 3a,b and Supplementary** Fig 4a-d). Pseudotime analysis showed bidirectional maturation trajectories from the d16 tuberal progenitors towards either neurons or tanycytes, indicating multipotential differentiation capacity of the early tuberal progenitors (**Fig. 3b-d**). Investigation of gene expression patterns along the tanycyte maturation trajectory revealed a gradual downregulation of *RAX* and *SOX2*, and transition through an intermediate *NOSTRIN+* progenitor state towards mature tanycytes expressing *CRYM, COL25A1, HTR2C and DIO2* together with the pan-glial markers *NFIA and NFIB* (**Fig. 3d,e**). Notably, whereas *SOX2*, *NFIA* and *NFIB* were also found to be expressed in human fetal astrocytes, *RAX*, *CRYM*, *DIO2*, *COL25A1* and *HT2RC* showed restricted expression to only the tanycyte population in the Herb *et al.* human fetal tissue dataset (**Supplementary** Fig. 4g,h). The mature tanycytes segregated into subpopulations expressing either *CRYM, GFPT1* or the serotonin receptor *HTR2C* (**Fig. 3e**). Through ICC, we confirmed the presence of tanycytic cells labelling positive for S100b and Vimentin but negative for the astrocytic marker AQP4 (**Supplementary** Fig. 4e). Stainings of four months old ARC grafts further confirmed the presence of human tanycytes indicated by the co-expression of human nuclear antigen (HuNu) with *RAX*, NFIA and Vimentin (**Fig. 3f**).

**Fig. 3.**
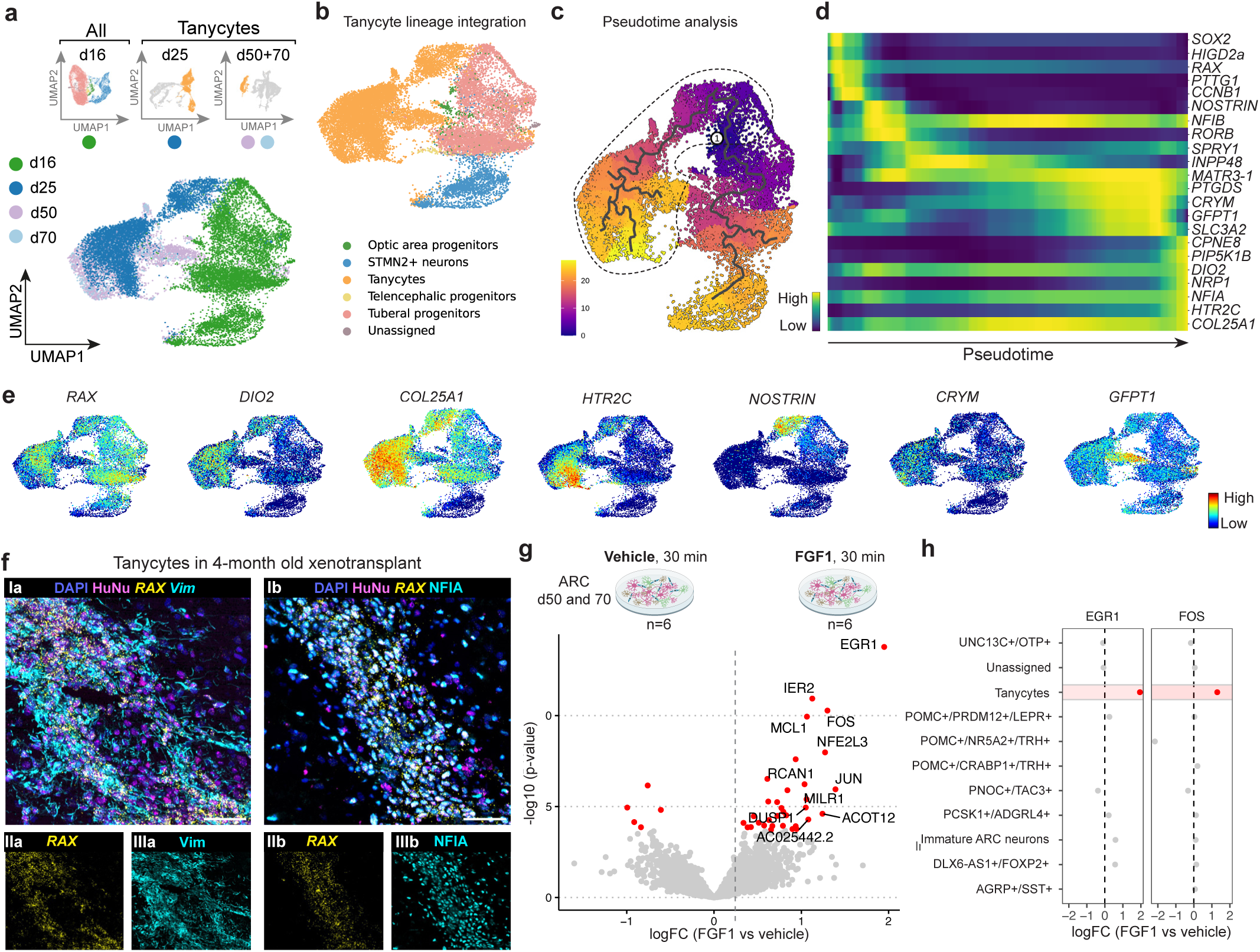
ARC protocol gives rise to mature tanycytes *in vitro* and in xenografts. **a**, Integration of day (d) 16 progenitor scRNAseq dataset with tanycyte cluster of d25, 50 and 70 snRNAseq datasets for analysis of the tanycyte lineage. **b,** Resulting integrated tanycyte dataset with annotated clusters. **c**, Pseudotime trajectory (Monocle 3) of integrated tanycyte dataset. (1) indicates pseudotime root. The dotted line encircles the tanycyte lineage used in d. **d**, Gene expression of key tanycyte and glial markers along the pseudotime trajectory. **e**, Feature plots of key markers expressed in the tanycyte dataset. **f**, Combinatorial immunofluorescence/*in situ* hybridization (IHC/ISH) of *RAX* (lIa and IIb), Vimentin (Vim; IIIa,), and NFIA (IIIb) within graft site (human nuclear antigen (HuNu) co-stain [Ia and Ib]). Scale bars, 50 µm. **g**, Top: Schematic overview of d50 and d70 arcuate nucleus (ARC) cultures treated with either vehicle or 50 ng/mL FGF1 for 30 min prior to nuclei extraction for snRNAseq (n=6 replicates per treatment, 3 from d50 and 3 from d70). Bottom: Volcano plot depicting differentially expressed genes in tanycyte cluster after stimulation. **h**, Differential expression (vehicle vs. FGF1 stimulation) of early response genes (*EGR1* and *FOS*) across all annotated cell types in the dataset showing significant upregulation in tanycytes in red.

Having gained direct access to human tanycytes and ARC neurons in the dish, we next investigated if the cultures could recapitulate ARC-mediated responses to FGF1; a growth factor which has been shown to mediate sustained remission of diabetic hyperglycaemia upon injection into the brain in mouse^32,33^. To this end, we treated d50 ARC cultures with FGF1 for 30 minutes and performed snRNAseq analysis of all cells in the culture (**Fig. 3g**). The treated cultures showed a FGF1 response exclusive to tanycytes as evidenced by strong induction of the early response genes *FOS* and *EGR1* in this cluster compared to the vehicle control (**Fig. 3h**). We thereby confirmed that the ARC protocol presented in this study produced fully mature and functionally responsive human tanycytes, originating from human tuberal progenitors.

### BMP timing for fine-tuning ARC fate

Previous data from the chick has uncovered that a BMP wave in the early embryo is responsible for patterning of the tuberal hypothalamus into an anterior POMC^+^/RAX^+^/SHH^+^ domain and a posterior FGF10^+^/TBX3^+^/RAX^+^ domain^27^ (**Fig. 4a**). Using our human stem cell model, we investigated whether this developmental phenomenon could be recapitulated *in vitro* through timed addition of BMP to the cultures between d3 and6 of differentiation (**Fig. 4b**). In our d16 progenitors, we showed that while *RAX* was highly expressed in all conditions, only the delayed addition of BMP4 at d5 or d6 induced high levels of posterior tuberal markers *TBX3* and *FGF10* while early BMP4 addition at d3 caused a profound patterning shift towards eye field as indicated by *PAX6* and *VSX2* expression (**Fig. 4c,d**). These findings were confirmed by combinatorial ICC/ISH, which also showed that late addition of BMP4 at d6 resulted in contaminating cell populations of telencephalic FOXG1^+^ phenotype (**Fig. 4e,f**). Comparison of mature d50 cultures derived from the BMP timing conditions showed that while all BMP-stimulated tuberal progenitors had the potential to generate POMC neurons, only the posterior fates stimulated with BMP from d5 or d6 could generate AGRP neurons (**Fig. 4g and Supplementary** Fig. 5a). Thereby, we conclude that progenitors with high *FGF10* and *TBX3* expression are required for the generation of AGRP neurons, and that initiation of BMP stimulation on d5 was most optimal for obtaining such cells without eye field and telencephalic contamination.

**Fig 4.**
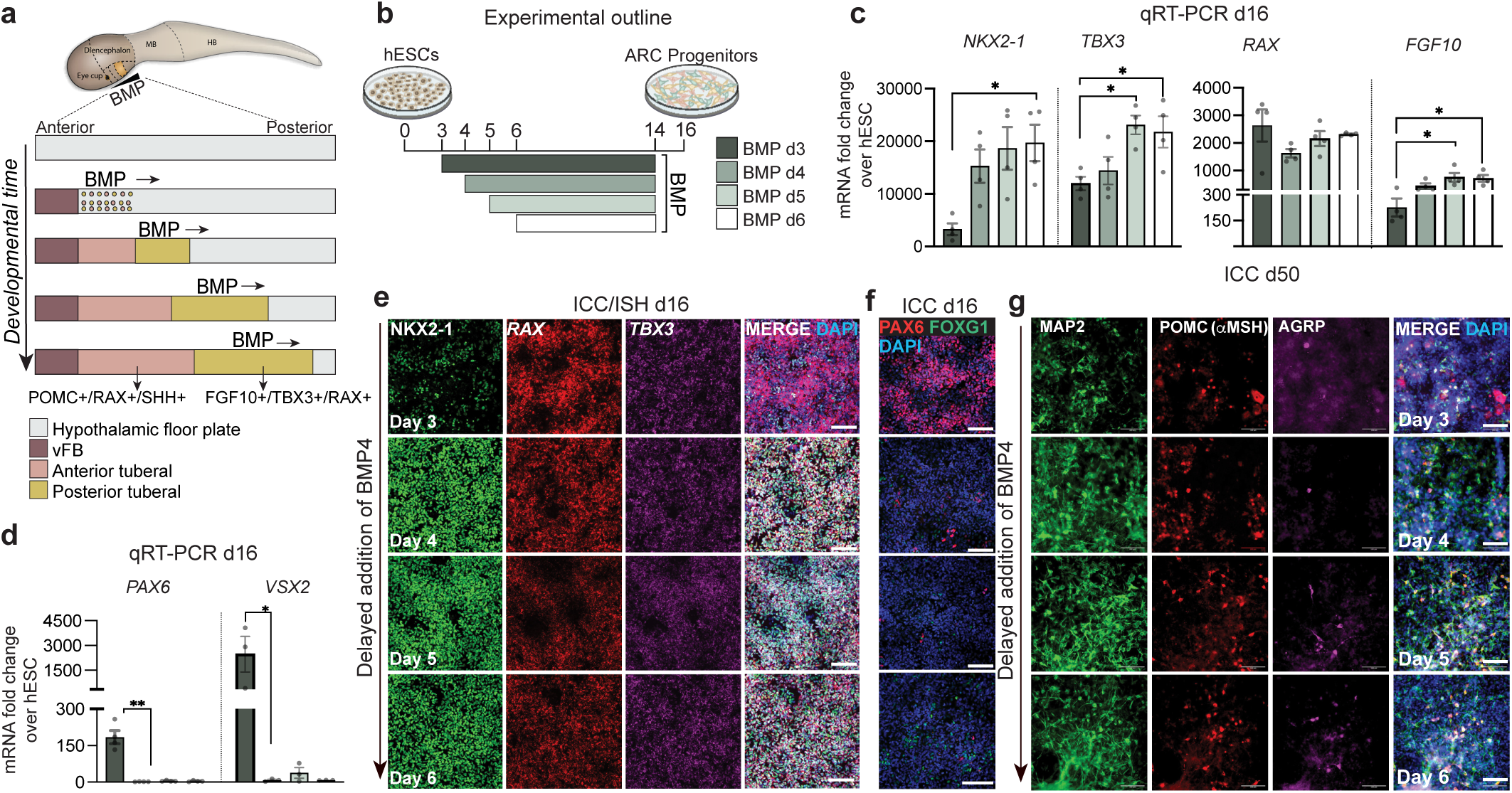
Early BMP addition induces eye field development at the expense of ARC neurons. **a**, Schematic representation of the BMP wave in the tuberal hypothalamus of the developing chick embryo specifying anterior (POMC+/SIX6+/ SHH+) and posterior (FGF10+/TBX3+/RAX+) in a time-dependent manner. Adapted from Chinnaiya *et al.*, 2023^21^. **b**, Schematic representation of the experimental design used to simulate BMP temporal patterning *in vitro*. BMP4 was added to cultures from either day (d) 3, 4, 5 or 6 until d14. **c-d**, qRT-PCR for early ARC (*NKX2-1*, *TBX3*, *RAX*, *FGF10*) and eye (*PAX6*, *VSX2*) markers from d16 ARC progenitors differentiated with BMP4 added on different starting days. Brown-Forsythe ANOVA with Dunnett’s T3 multiple comparisons test: *NKX2-1*: DAY3vsDAY5 *p= 0.0443, DAY3vsDAY6 *p= 0.0141. One-way ANOVA with Tukey’s multiple comparisons test: *TBX3*: DAY3vsDAY5 *p= 0.0208, DAY3vsDAY6 *p=0.0432; *FGF10*: DAY3vsDAY5 p=0.0293, DAY3vsDAY6 *p=0.0437. Kruskal-Wallis with Dunn’s multiple comparisons test: *PAX6*: DAY3vsDAY4 **p= 0.0050; *VSX2*: DAY3vsDAY4 *p= 0.0140. **e**, Combinatorial immunocytochemistry (ICC) (NKX2-1) and *in situ* hybridisation (ISH) (*RAX, TBX3*) on d16 ARC cultures treated with BMP4 starting at different start days. Scale bar, 100 µm. **f**, ICC of FOXG1 and PAX6 in d16 ARC progenitors treated with BMP4 starting at different time points. Scale bar, 100 µm. **g**, ICC of d50 ARC cultures for POMC(αMSH), AGRP, and MAP2. Scale bar: 100 µm. ARC, arcuate nucleus; BMP, bone morphogenetic protein; FB, forebrain; HB, hindbrain; MB, midbrain; vFB, ventral forebrain.

BMP7, in addition to BMP4, has been equally implicated in hypothalamic development in animal models^20^. Thus, we investigated if BMP7 could also induce early tuberal ARC specification. We found no significant difference in the expression of NKX2-1, *RAX, TBX3, ISL1* and *FGF10* when substituting BMP4 with BMP7, suggesting that these two factors can act redundantly and with equal efficiency in the specification of early tuberal fates (**Supplementary** Fig. 5b,c). Collectively, these data show that it is the timing rather than the subtype of BMP which is crucial for correct posterior tuberal induction and generation of AGRP neurons *in vitro*.

### Early BMP withdrawal generates VMH

Further investigating the BMP wave hypothesis (**Fig. 4a**), we reasoned that the cells might be affected not only by the onset but also by the duration of BMP signalling. A simple way to test this *in vitro* was to add BMP at d5, and then withdraw it again at different timepoints between d7-14, including a control condition without BMP (**Fig. 5a**). Combined ICC/ISH of the progenitors at d16 showed that only conditions where BMP was kept in the medium until d11 or d14 displayed high expression of *TBX3*, which is a downstream target of BMP (**Fig. 5b,c**). In contrast, earlier withdrawal of BMP4 at d7 or d9 caused a gradual decrease in TBX3 and *RAX* expression levels (**Fig. 5c** and **Supplementary** Fig. 6a). The posterior marker *FGF10* was also significantly reduced in the shorter BMP treatment conditions, while the anterior marker *SHH* was upregulated (**Fig. 5b**). Analysis of the same BMP withdrawal conditions at d50 showed that while the posterior progenitors receiving BMP4 from d5-11 or d5-14 were capable of generating ARC-related AGRP and TRH neurons, the anterior progenitors receiving shorter BMP stimulation (d5-7 or d5-9) were largely devoid of these neuronal subtypes (**Fig. 5d,e and Supplementary** Fig. 6b,c). We further found that the longest BMP4 treatment (d5-14), generating the most posterior tuberal fates, induced the highest expression of ARC tanycyte markers *DIO2* and *RAX* (**Fig. 5d**). In contrast, the anterior conditions displayed characteristic features of the neighbouring VMH nucleus as evidenced by the expression of VMH markers *SOX14*, *GPR149* and NR5A1 (**Fig. 5d,e**). In contrast, *PRDM12* and *POMC*/αMSH expression was found in a broader range of BMP conditions, including also in the condition where no BMP was added (**Fig. 5e and Supplementary** Fig. 6b), indicating a broader developmental origin of POMC-expressing neurons. To investigate robustness of the findings across both hESC and human induced pluripotent stem cell (hiPSC) lines, we replicated the BMP withdrawal experiment in a widely used induced pluripotent stem cell (hiPSC) line, KOLF2.1. This data from a separate hPSC line confirmed that only posterior tuberal progenitors generated by prolonged BMP stimulation had the capacity to generate AGRP neurons whereas shorter BMP treatment (d5-9) generated NR5A1^+^ VMH cells (**Fig. 5f and Supplementary** Fig. 6b-d).

**Fig 5:**
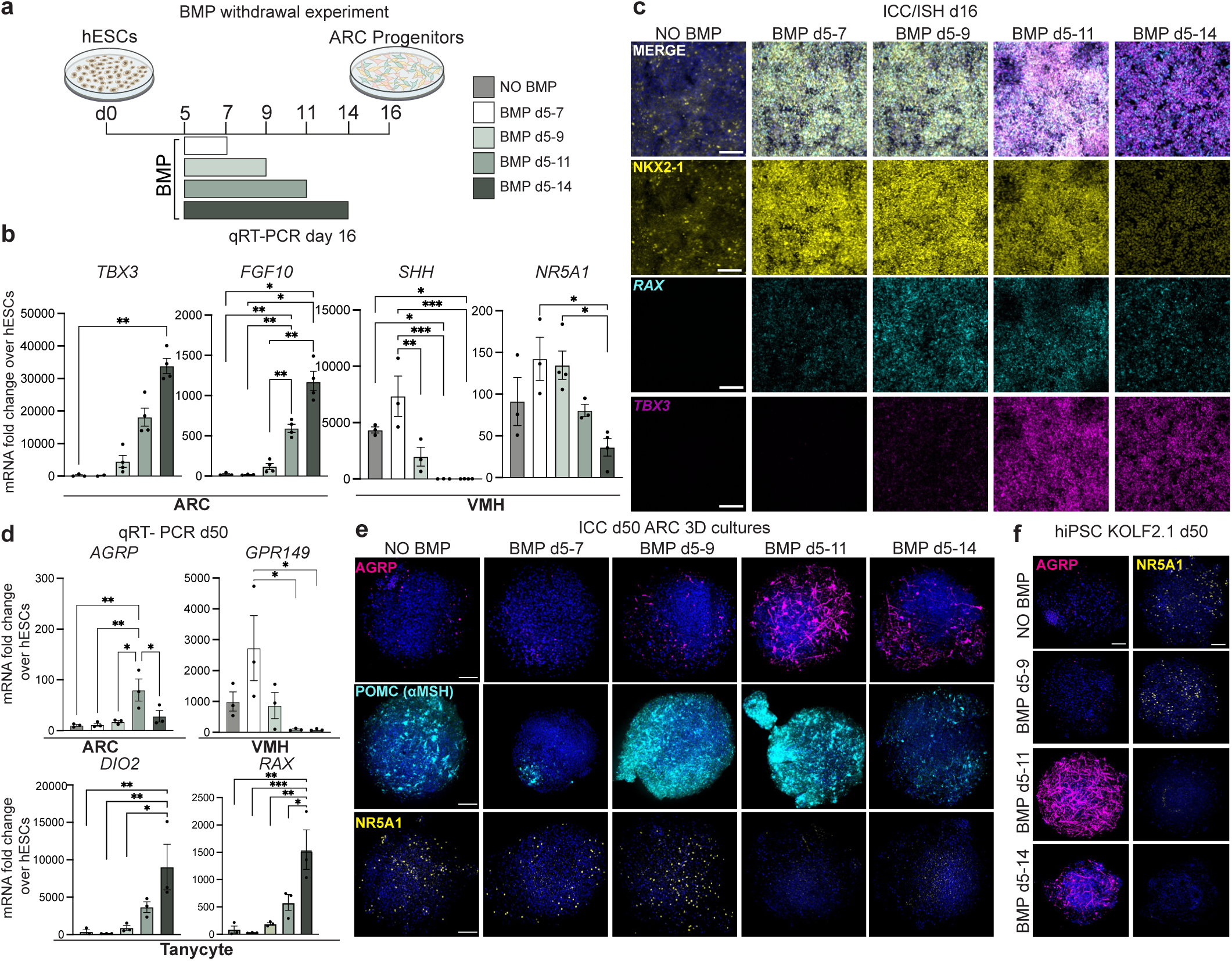
BMP from d5-11 is ideal for generation of posterior tuberal-derived AGRP neurons. **a**, Schematic representation of the experimental design used to simulate BMP temporal withdrawal *in vitro*. BMP4 was added to cultures from day (d) 5 to either d7, 9, 11, or 14 or omitted (neg. control). **b**, qRT-PCR of early posterior (*TBX3, FGF10*) and anterior (*SHH, NR5A1*) tuberal markers in d16 ARC progenitors differentiated with different termination days of BMP4 treatment. Kruskal-Wallis with Dunn’s multiple comparisons test: *TBX3*: noBMPvsd5-14 **p=0.0088. Brown-Forsythe ANOVA with Dunnett’s T3 multiple comparisons test: *FGF10*: noBMPvsd5-14 *p= 0.0125, d5-7vsd5-14 *p= 0.0122, noBMPvsd5-11 **p= 0.0079, d5-9vs5-14 **p= 0.0069, d5-9vsd5-11 **p= 0.0021, d5-7vsd5-11 **p= 0.0072. One-way ANOVA with Tukey’s multiple comparisons test: *SHH*: noBMPvsd5-14 *p=0.0180, d5-7vsd5-14 ***p=0.0003, noBMPvsd5-11 *p=0.0272, d5-7vsd5-11 ***p=0.0006, d5-7vsd5-9 **p=0.0068; *NR5A1*: d5-7vsd5-14 *p=0.0115, d5-9vsd5-14 *p=0.0112. **c,** Combinatorial immunocytochemistry (ICC) (NKX2-1) and in situ hybridisation (ISH) (*RAX*, *TBX3*) on d16 ARC cultures with different days of BMP4 withdrawal. Scale bars, 100 µm. **d**, qRT-PCR of d50 ARC cultures with markers for ARC (*AGRP*), VMH (*GPR149*), and tanycytes (*DIO2*, *RAX*). One-way ANOVA with Tukey’s multiple comparisons test: *AGRP*: noBMPvsd5-11 **p=0.0086, d5-7vsd5-11 **p=0.0098, d5-9vsd5-11 *p=0.0173, d5-11vsd5-14 *p=0.0493; *GPR149*: d5-7vsd5-14 *p=0.0335, d5-7vsd5-11 *p=0.0340; *DIO2*: noBMPvsd5-14 **p=0.0100, d5-7vsd5-14 **p=0.0084, d5-9vsd5-14 *p=0.0150; *RAX*: noBMPvsd5-14 **p=0.0012, d5-7vsd5-14 ***p=0.0009, d5-9vsd5-14 **p=0.0021, d5-11vsd5-14 *p=0.207. **e**, ICC of AGRP, POMC(αMSH), and VMH marker NR5A1 in d50 ARC 3D cultures of BMP withdrawal experiment. Scale bars, 100 µm. **f**, BMP withdrawal experiment performed using a human induced pluripotent stem cell (hiPSC) line (KOLF2.1). ICC of ARC marker, AGRP, and VMH marker, NR5A1, in d50 ARC 3D cultures. Scale bars, 100 µm.

In summary, we show that by manipulating the timing and duration of BMP stimulation, we could mimic the anterior-to-posterior BMP wave that has been shown to pattern distinct areas along the hypothalamic floorplate in the tuberal hypothalamus of the chick^27^. Through this approach, we were able to control subregional patterning of human pluripotent cells into either anterior SHH^+^/SOX14^+^/NR5A1^+^ VMH progenitors or posterior RAX^+^/TBX3^+^/FGF10^+^ ARC progenitors giving rise to AGRP neurons. Across two different hPSC lines, BMP stimulation from d5-11 proved to be the ideal condition for generating ARC cultures enriched for AGRP neurons. In line with what has been suggested from the chick^27^, our data further suggests that human tanycytes arise from posterior FGF10^+^ tuberal progenitors, potentially even more posterior than the progenitors giving rise to AGRP neurons (**Fig. 6**).

**Fig 6.**
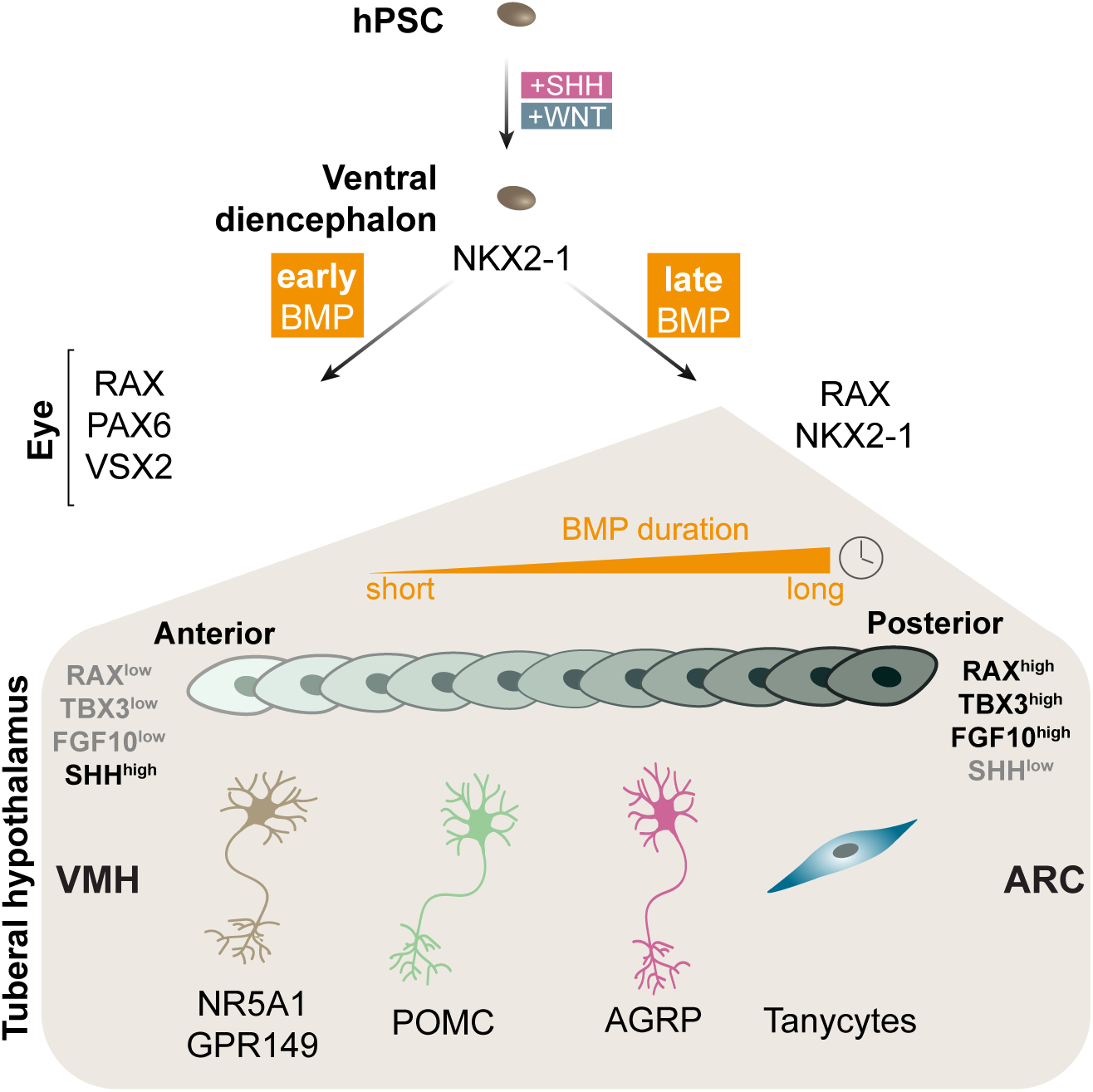
Anteroposterior patterning of the tuberal hypothalamus is dependent on BMP. WNT and SHH activation together with dual SMAD inhibition for neural induction patterns human pluripotent stem cells (hPSC) towards ventral diencephalic fate (NKX2-1+). Early onset of BMP induces optic fate (PAX6, VSX2, RAX) whereas later BMP generates NKX2-1+/RAX+ hypotha-lamic progenitors. These tuberal hypothalamic progenitors can be patterned towards an anterior versus posterior fate depending on the duration of BMP exposure. Early withdrawal of BMP generates anterior progenitors with high expression of SHH and low expression of RAX/TBX3/FGF10. In contrary, longer exposure to BMP induces high RAX, TBX3 and FGF10 expression and downregulates SHH. The most anterior tuberal progenitors give rise to VMH populations defined by NR5A1 and GPR149. Within the ARC-committed cells, the more anterior populations generate high numbers of POMC neurons whereas AGRP neurons and tanycytrs can only develop from the more posterior regions with long BMP exposure.

## Discussion

The generation of stem cell-derived *in vitro* models of the hypothalamus has been challenged by a lack of knowledge on subregional lineage trajectories and of specific markers identifying lineage-committed progenitors. While a few hypothalamic lineages have been tracked through lineage fate mapping^34,35^, the developmental origin of most hypothalamic subtypes is still unaccounted for, and speculative presumptions on progenitor-to-neuron lineages have been based mainly on anatomical gene expression patterns. *In vitro* stem cell models can fill this gap by allowing longitudinal access to individual lineages without the confounding effects of anatomical folding and inter-regional migration that can occur in the hypothalamus during development^36,37^. Through data-driven protocol optimisation we show here that we could successfully optimise protocols towards PVN versus ARC fates, an approach which is applicable to generate future protocols towards other regions of the hypothalamus in a similar manner. This data showed us that a posterior tuberal combinatorial expression profile of NKX2-1, TBX3 and RAX was required for the derivation of key ARC lineages, including AGRP, PNOC and tanycyte subtypes. Whilst previous differentiation protocols have yielded high purity of NKX2-1 expressing progenitors, they have been heterogenous in their sub-regional fates with the highest reported purity being 35% for RAX^17^ and 10% for TBX3^16^. In line with our findings showing a broader developmental origin of POMC-expressing neurons, these previous protocols have generated mainly POMC neurons, but lacked the presence of other important ARC-derived cell types such as AGRP and PNOC neurons, as well as tanycytes.

Here, we could generate such specific ARC cell types through both in vitro and in vivo xenograft maturation of lineage-committed d16 posterior tuberal progenitors, and we further validated the transcriptional profile of the mature neurons and tanycytes against human arcuate nucleus snRNAseq data. We identified several subtypes of human POMC neurons, including two canonical POMC clusters, namely POMC^+^/NR5A2^+^/TRH^+^ and POMC^+^/PRDM12^+^/LEPR^+^, enriched in 3D and 2D, respectively. Both subtypes expressed high levels of *PRDM12* and *LEPR*, likely corresponding to the well-characterised murine POMC neurons known to regulate energy and blood glucose homeostasis^2,38,39^. We further validated our model by showing that a subset of the POMC^+^/LEPR^+^/PRDM12^+^ population also co-expressed *NR5A2*, consistent with data from the mouse^40^. Moreover, we found multiple ARC populations expressing *PNOC*, including the GHRH^+^/PNOC^+^ cluster as well as a PNOC^+^/TAC3^+^ cluster. These ARC PNOC-expressing neurons showed similarities to a recently discovered GABAergic PNOC population in the mouse, which has been shown to promote food intake when activated^29,30,41^.

The derivation of the human ARC lineage was highly dependent on the timing of BMP stimulation, comparable to the anterior-to-posterior BMP wave which has been identified in the chick tuberal hypothalamus^27^. Whilst sequential addition of BMP has been employed to study neural tube patterning of the dorsoventral axis in an *in vitro* mouse ESC differentiation system^42^, its application in human ARC-oriented differentiation has not previously been explored. As in the chick^27,43^, we found that the induction of *TBX3* by BMP was essential for downregulation of *SHH* and upregulation of *FGF10* to obtain correctly patterned posterior tuberal hypothalamic fates. While the study in the chick did not uncover the resulting adult hypothalamic nuclei or neuronal subtypes arising from each tuberal progenitor domain, our human *in vitro* system demonstrates that the anterior to posterior tuberal progenitor domains are likely to underlie the developmental origins of the VMH and the ARC, respectively. We further show that it is possible to fine-tune *in vitro* patterning with BMP timing to a degree where we could enrich the ARC cultures for either AGRP or tanycyte lineages, indicating that tanycytes are derived from a tuberal domain which is posterior to that giving rise to AGRP neurons. These findings corroborate indicative data from the chick showing that FGF10^+^ posterior tuberal progenitors at late developmental stages upregulate tanycytic markers^27^. Although the *in vitro* derived tanycytes expressed several mature tanycyte markers, we were not able to identify equivalents to the α or β tanycyte subtypes that have been identified in the mouse^44^. It is conceivable that region-specific spatial cues along the 3^rd^ ventricle *in vivo* are required to obtain distinct α and β tanycytic identity. Also, tanycyte subclassification has not been transcriptomically characterised in the human, so it is uncertain if subtype-specific markers are conserved between mouse and human. We further leveraged our cultures for functional testing and provided evidence from a human perspective to support previous data from the mouse identifying tanycytes as the primary responders to FGF1 in the adult hypothalamus^33,45^.

In sum, our novel protocol and proof-of-concept functional experiments provide a foundation for efficient screening of molecular effects of biological and pharmacological molecules on human subtype-specific hypothalamic functions. We have developed that system by applying developmental insights from model organisms to fine-tune protocols for the directed subregional differentiation of hypothalamic fates from human stem cells. Access to *in vitro* models of this type will enable detailed investigation of the cellular and molecular responses to relevant nutrients, hormones and drug candidates, as well as CRISPR-based perturbation screens of obesity-associated genetic loci, to identify and characterise hypothalamic energy homeostatic pathways in humans.

## Reporting summary

Further information on research design is available in the Nature Portfolio Reporting Summary linked to this article.

## Acknowledgements

We want to especially thank Helle Lilja-Fischer, Kristoffer L. Egerod and the Single-Cell Omics platform at the Novo Nordisk Foundation Center for Basic Metabolic Research (CBMR) for technical support on scRNAseq and snRNAseq experiments, the Microscopy and Genomics platforms at reNEW Copenhagen and Arun Thiruvalluvan for help with imaging, Jette Pia Larsen and Sonia Araujo for technical assistance, Lorenzo Fedrizzi for cell culture maintenance and Gaurav Rathore for advice on bioinformatic analysis. This study was supported by the Danish Research Council (grant no. 0169-00073B), the European Union (H2020, NSC-Reconstruct GA no 874758), the Lundbeck Foundation (R350-2020-963 and R380-2021-1267) and the Knut and Alice Wallenberg Foundation. The Novo Nordisk Foundation Center for Stem Cell Medicine (reNEW) and Center for Basic Metabolic Research (CBMR) are supported by donations from the Novo Nordisk Foundation (Grant number NNF21CC0073729 and NNF23SA0084103).

## Author contributions

AK, ZA, AKM and EH conceived the study. ZA, AKM, JBC, VN and SP performed the *in vitro* differentiations and downstream analyses. EH, DR and YL performed all bioinformatic analysis. AS and ALS performed *in vivo* transplantation of cells and LP did analysis of rat brain sections. JK performed the calcium imaging. ZA, AKM, SP, JBC collected the data and AK, AKM, ZA and EH wrote the manuscript with input from DR and THP.

## Ethics declaration

This study uses previously derived hESC and hiPSC cell lines RC17 and KOLF2.1, respectively. The RC17 and KOLF2.1 cell lines have been derived under local ethical approval at the original derivation sites and with informed consent from donors. No new hESC or hiPSC lines were derived in this study. The Kirkeby lab holds a relevant ethical approval to work with hPSC lines (H-21043866). Xenotransplantation studies to rats were approved by a local animal research committee and performed under animal ethics license 2022-15-0201-01236.

## Methods

### Cell culturing and neuronal differentiation

One hESC line called RC17 (Roslin Cells, hPSCreg: RCe021-A) and one hiPSC lines called KOLF2.1J^1^ were used in this study. Stem cells were maintained in IPS-Brew (Miltenyi, #130-104-368) on plates coated with 1 μg/cm^2^ Laminin-521 (Biolamina, #LN521-02) in phosphate-buffered saline with Mg^2+^ + Ca^2+^ (PBS+/+). Cells were passaged at 70-90% confluency with 75 μL/cm^2^ 0.5 mM EDTA (Thermo Fisher Scientific, #15575020) for 5-7 min, then detached and replated in IPS-Brew. Medium was supplemented with 10 μM Y-27632 (ROCK inhibitor [ROCKi], Miltenyi, #130-106-538) for 24 hours with daily media changes thereafter. For differentiation into regionalized hypothalamic progenitors, we adapted the protocol from Nolbrant *et al.*, 2017^2^. All cultures received N2 medium from d0 to 11 (50% MACS Neuromedium, 50% DMEM/F12 containing 1% N-2 supplement [Thermo Fisher Scientific, #17502048] 1% GlutaMAX [Thermo Fisher Scientific, #35050061], 10 U/mL Penicillin/Streptomycin (Gibco, #15070063) and NB21 maturation medium from d11 until end of experiment (MACS Neuromedium containing 2% NB-21 supplement, 1% GlutaMAX, Penicillin/Streptomycin]. Medium was changed every 2-3 days and whenever cells were replated, 10 µM ROCKi was supplemented. At d0, hESC were dissociated as described, washed in DMEM/F12 with 5% KOSR (Gibco, #10828-028) and plated at 10K/cm^2^ in plates coated with 2 μg/cm^2^ laminin-111 (Biolamina, #LN111-02) in PBS+/+. At d11, progenitors were dissociated with accutase (Thermo Fisher Scientific, #A11105-01) for 10 min and replated at 800k/cm^2^ onto plates coated with 2 μg/cm^2^ laminin-111 in PBS+/+. After the last replating at d16, cells were replated at 700k/cm^2^ onto 2 μg/cm^2^ laminin-521 in PBS +/+ coated plates. All differentiations received 10 μM SB413542 (Miltenyi, #130-106-543), and 100 ng/mL Noggin (Miltenyi, #130-103-456) in N2 basal medium from day 0 to 9, 0.2 mM ascorbic acid (Sigma-Merck, #A4403-100MG) and 20 μg/mL BDNF (Miltenyi, #130-096-286) from d11 and 500 μM dibutyryl-cAMP (Sigma-Merck, #D0627-1G), 1 μM DAPT (Miltenyi, #130-110-489), 10 ng/mL GDNF (Miltenyi, #130-098-449) from d16 onwards. The NB21 medium supplemented with factors (AA, BDNF, db-cAMP, DAPT, GDNF) from d16 was named maturation medium (MM). For the PVN protocol, 0.3 µM CHIR was added day 0 to 9 and 150ng/ml SHH from day 6 to 14. The ARC protocol included 0.3 µM CHIR d2-9, 300 ng/mL SHH day 0 to 9 and 400 ng/mL IGFBP-3 (R&D systems, #675-B3-025) from day 7 to 14. In addition, 50 ng/mL BMP4 (Milteny, #130-111-168) was supplemented at varying timepoints. In the ARC protocols, cells received 100 ng/mL BMP-7 (R&D systems, #354-BP-500/CF) and 4mM dimethyl 2-oxoglutarate (dm-aKG, Sigma, #349631) from day 25 to 50 of maturation.

For the generation for spheroids, patterned d16 hypothalamic progenitors were thawed and diluted in pre-warmed DMEM/F12 with 5% KOSR. Cells were counted, centrifuged at 400 xg for 5 min and resuspended in NB21 medium with 20 ng/mL BDNF, 0.2 mM ascorbic acid, 500 µM Db-cAMP and 1 µM DAPT at 250 000 cells/mL. The cell suspension was transferred to an ultralow attachment U-bottom 96-well plate (Costar) at 200 µL/well, centrifuged at 100 xg for 5 min and maintained at 37 °C with 5% CO_2_. 75% of maturation medium was changed every 2-3 days until termination of the experiment.

### Xenotransplantation of day 16 ARC progenitors

All procedures were conducted in accordance with the European Union Directive (2010/63/EU) and had approval by the local ethical committee at Lund University and the Swedish Department for Agriculture (Jordbruksverket). Seven adult, female, athymic nude rats (Hsd:RH-Foxn1^rnu^, Inotiv (Prev. Envigo), Indiana, USA) were housed on a 12:12-hr light:dark cycle with *ad libitum* access to food and water. ARC cells were prepared for transplantation by thawing day 16 cryopreserved progenitors in wash buffer (0.5% human serum albumin (HSA) in HBSS-/-) and centrifuged at 400 xg at RT. Cells were reconstituted to 167,000 cells/µL in Neurobasal (Thermo Fisher Scientific, #21103049) with 20 U/mL pulmozyme, 1x B27 supplement (Thermo Fisher Scientific, #17504044), 10 µM ROCKi, 0.2 mM ascorbic acid and 20 ng/mL BDNF.

For unilateral intrastriatal xenotransplantations, the rats (>225 g) underwent general anesthesia (Fentanyl (45 mg kg^-1^)-Domitor (0.03 mg kg^-1^) mix, Apoteksbolaget, i.p.). A glass capillary was inserted into the striatum at the following coordinates for two deposits: AP_1_: +0.9, ML_1_: -3.0, DV_1_: -5.0; AP_2_: +1.4, ML_2_: -2.6, DV_2_: -5.0. The cells were infused in a volume of 0.1 µL per 12 s over 3 min (1.5 µL of 250,000 cells in total) followed by 2 min diffusion time per deposit.

### RNA extraction and qRT-PCR (quantitative real time PCR)

Approx. 300-500K cells were lysed using 350 μL of RLT buffer (QIAGEN, #74034) containing 0.5 mM beta-mercaptoethanol (Thermo Fisher Scientific, #31350010), snap frozen and stored at -80°C. RNA isolation was performed with a QIAcube using the RNeasy plus micro kit (Qiagen, #74034). Approx. 1ug of RNA was reverse transcribed using the Maxima first strand cDNA synthesis kit (Thermo Fisher Scientific, #K1642). cDNA was diluted in 250ul EB buffer (Qiagen, #19086) and stored at -20°C degrees. For qRT-PCRs, SYBR green (Roche, #04887352001), primers (see sequences in **Supplementary Table 1**) and cDNA sample were pipetted by an iDOT liquid handler and run on a Light Cycler 480 II instrument (Roche, #05015243001) with 40 cycles-60°C for 60 s annealing/elongation step and 95°C, 30s denaturation. The average CT of technical duplicates was used to calculate the relative expression and this was normalized to a reference consisting of the average expression of undifferentiated H9 and RC17 cells. Genes of interest were normalized to both *GAPDH* and *ACTB*. Resulting change in fluorescence intensity was plotted in Prism (GraphPad v10.2.3).

### Analysis of hypothalamic expression matrix from qPCR data

qRT-PCR expression data from early differentiations were log-normalized prior to principal component analysis. To identify the effects of distinct factors on PC1 embedding, a linear model was constructed to predict PC1 embedding values, including all tested compounds. The resulting model was analyzed using estimated marginal means to identify the average effects of different factors while adjusting for other covariates in the model. To identify genes in early differentiations which are predictive of marker genes in late differentiations, a linear model was constructed to test the relationship between each individual gene and the selected late marker. Gene expression values were log normalized and scaled prior to analysis. Only differentiations in which the two genes had been measured at the early and late timepoints were included.

### Immunocytochemistry (ICC)

All 2D cultured cells were fixed in 4% paraformaldehyde for 15 min at 37°C followed by three washes with phosphate-buffered saline (PBS-/-) and stored at 4°C until staining. Cells were blocked for 1-3 h at room temperature (RT) in blocking buffer (0.1%, Triton X-100 and 5% (vol/vol) donkey serum in PBS-/-). Primary antibodies (see **Supplementary Table 2**) were diluted in blocking buffer and incubated at 4°C overnight. Following incubation, cells were washed three times with PBS-/-before incubation with secondary conjugated fluorophores (1:200, see **Supplementary Table 3**) and 2 µg/mL DAPI for 2 h at RT on a shaker. A final three PBS washes were performed before imaging.

Spheroids were fixed with 4% PFA for 30 min at RT, washed twice with PBS and blocked for 6 h at 4°C with blocking buffer. Thereafter, they were incubated with primary antibodies diluted in blocking buffer (200µL/tube) on a shaker at 4°C for 48 h. The primary antibody solution was removed and the samples were washed with 300 µL blocking buffer, once for 2 min and then for 6 h at 4°C. All subsequent steps were performed under the exclusion of light. The secondary antibodies and 2 µg/mL DAPI for nuclear counterstaining were diluted in 200 µL blocking buffer and added to the tissues. The samples were again incubated on a shaker at 4°C for 48 h. Subsequently, the tissues were washed for 2 min and then at 4°C for 6 h on a shaker. After two more washes in PBS, the tissues were transferred to 18-well ibidi chambers and stored at 4°C until visualization.

### Multiplexed immunohistochemistry

The nude rats were anaesthetized with Sodium Pentobarbital (250-350 mg/kg, 1.4 mL of 60 mg/mL per rat, Apotek, Sweden) and transcardially perfused with 0.1 M phosphate buffer. The brains were removed, post-fixed in 4% PFA for 24 h at RT and immersed in 25% sucrose solution until fully dehydrated. For sectioning, the brains were cut into 35 µM coronal sections using a cryostat (Leica Microsystems, CM1950 cryostat) and systematically sampled in series of eight. The sections were stored in cryoprotectant at -20 °C until use. Sections encompassing the striatum (site of xenotransplant), and the ARC/PVH (endogenous positive controls) were used for immunostainings.

For immunostainings, the sections were initially rinsed in PBS-/- and subjected to antigen retrieval in pre-heated (80°C) TRIS-EDTA buffer (10 mM Tris Base, 1 mM EDTA Solution, 0.05% Tween-20, pH 9) for 30 min, and left to cool to RT before rinsing in PBS-/-. For SG chromogenic staining, endogenous peroxidase activity was blocked by incubation in 1% hydrogen peroxidase (H_2_O_2_) in PBS-/- for 20 min at RT, prior to pre-incubation in blocking buffer (5% donkey or goat serum, 1% bovine serum albumin, 0.3% Triton X-100 (TX) (0.1% TX for SG chromogenic staining) in PBS-/-) for 30 min at RT. Next, the sections were incubated overnight (2 h for anti-hNCAM) at RT in primary antibodies diluted in blocking buffer. After primary antibody incubation, the sections were rinsed in washing buffer (0.25% bovine serum albumin, 0.1% Triton X-100 in PBS) or in 0.1% TX-PBS for SG chromogenic staining, followed by incubation for 1 h at RT in fluorophore-conjugated or biotin-coupled secondary antibodies diluted in washing buffer. See **Supplementary Table 2 and 3** for information on primary and secondary antibodies, respectively.

For immunofluorescence, the sections were thereafter rinsed once in washing buffer and twice in PBS-/-, mounted on glass slides, and coverslipped using FluorSave^TM^ (Millipore, #345789). For SG chromogenic staining, the sections were rinsed in 0.1% TX-PBS and incubated for 1 h at RT in avidin-biotin complex solution (Vector Laboratories, #AK-5000) diluted in TX-PBS according to manufacturer’s instructions. After incubation, the sections were rinsed once in TX-PBS and twice in PBS-/-before color development for 10 min in ImmPACT Vector SG Substrate chromogen solution (Vector Laboratories, #AK-4705) without H_2_O_2_, and for additional 2 min with H_2_O_2_. The sections were next mounted on gelatin-coated microscope glass slides, dehydrated in a series of ethanol (70%, 96%, 99%), cleared in xylene, immediately cover slipped using DPX New mountant (Merck, #1.00579), and left to dry overnight.

### In situ hybridization (ISH) assay on cell cultures

Cells were fixed for 30 min at RT. *In situ* hybridization was performed using the RNAscope® Multiplex Fluorescent Kit v2 (ACD BioTechne, #323100) according to manufacturer’s instructions for cultured adherent cells in 96-well plates. The probes and fluorophores used in this study can be found in **Supplementary Table 4**. For combinatory ICC/ISH, the standard ICC protocol was performed after the RNAscope protocol.

### In situ Hybridization (ISH)/immunofluorescence (IF) on brain sections

Rats were deeply anaesthetized with CO_2_ gas and sacrificed by decapitation. The brain was quickly removed and immediately immersed in dry ice and stored at -80°C until sectioning. The brains were sectioned into 12 µM coronal sections using a cryostat (Leica, CM1950 cryostat) and directly mounted onto SuperFrost® Plus glass slides (Thermo Scientific, #J1800AMNZ) and stored at -80°C. ISH was performed according to the RNAscope® Multiplex Fluorescent v2 Assay combined with Immunofluorescence (Doc. No. MK51-150 Rev B/Appendix C-D) with several changes. The sections were first fixed in pre-chilled 4% PFA at 4°C overnight prior to dehydration and incubation in hydrogen peroxide (ACD Bio-Techne, #322330 or 3% H_2_O_2_ in RNAse-free water). Next, the sections were subjected to co-detection target retrieval (#323166, ACD Bio-Techne, Milan, Italy) for 5 min at 98-100°C, rinsed in ddH_2_O, and transferred to 100% ethanol for 3 min. The sections were then incubated in primary antibodies at 4°C overnight. According to protocol, the sections were rinsed post-fixed and subjected to protease treatment (RNAscope^®^ Protease Plus (ACD Bio-Techne, #322330), 1:5 diluted in RNAse-free water) for 15 min at 40°C. Next, RNA-specific oligonucleotide target probes were hybridized, amplified, and developed using Opal Dye fluorophore 570. Following the RNAscope protocol, the sections were rinsed and incubated in secondary antibodies for 60 min at RT. Lastly, the sections were rinsed and coverslipped using VectaShield Vibrance antifade Mounting medium with DAPI (#H-1800, Vector Laboratories, CA, USA). See **Supplementary Table 2-4** for information on primary and secondary antibodies, probes and fluorophores.

### Imaging and processing

Imaging of ICC of *in vitro* derived cells was performed on the Leica AF600 widefield epifluorescence microscope (Plan-Fluotar 20x/0.40 Dry) and for the main figures, the confocal microscope Leica Stellaris 5 (Plan-apochromat 40x/1.25 GLYC). All imaging of 3D tissue staining was performed using either the Zeiss LSM880 (EC Plan-Neofluar 20x0.50 WD 2.0mm and C-Plan-Apochromat 63x1.40 Oil UV-IR WD 0.14mm) or an inverted confocal microscope (ECLIPSE Ti2-E, Nikon with CFI Plan Apochromat Lambda 20X/0.75, WD 1.0mm) equipped with a spinning disk module (CSU-W1, Yokogawa). Images were processed using ImageJ (version 2.1.0).

Images of xenotransplantation sections (single and stacks) were acquired by confocal fluorescence (Leica Stellaris 5, Plan-Fluotar 10x/0.30 Dry or Plan-apochromat 40x/1.25 GLYC), widefield fluorescence (AF6000, Plan-Fluotar 10x/0.30 Dry or Plan-Fluotar 20x/0.40 Dry) or widefield brightfield microscopy (Leica DM5500, plan-apochromatic 10x/0.40, DFC450 color camera). The images were processed using ImageJ and brightness/contrast were adjusted using Adobe Photoshop 2024.

### Calcium Imaging

Before the start of the experiment, cells cultured in 18-well µ-Slide (ibidi, #81826) were incubated for 30 min with 3 µM Calbryte 520 AM calcium indicator (AAT Bioquest, #20650) in BrainPhys Imaging Optimized Medium (STEMCELL Technologies, #5796) with 0.02% Pluronic F-127 (Sigma Aldrich) for 30 min at 37°C. Cells were then rinsed with BrainPhys Imaging Optimized Medium and transferred to an inverted confocal microscope (ECLIPSE Ti2-E, Nikon) equipped with a spinning disk module (CSU-W1, Yokogawa) and environmental chamber. 40x water immersion objective (N.A. 1.25) was used for imaging. For potassium chloride (KCl) stimulation, cells were continuously imaged at 5 Hz. After 5 seconds, KCl (SigmaAldrich) at 50 mM final concentration was added to the well during imaging and corresponding response recorded for 2 minutes. For leptin stimulation experiment, baseline images were acquired first and then half of the medium in the well was replaced with the stimulant (leptin in BrainPhys Imaging Optimized Medium at 100 ng/ml final concentration) or vehicle (BrainPhys Imaging Optimized Medium). A second set of images was acquired after 30 min. Images were analyzed in ImageJ where mean fluorescence intensity per cell was extracted before and after stimulation. Resulting change in fluorescence intensity was plotted in Prism (GraphPad v10.2.3).

### AGRP ELISA

To measure the concentration of secreted AGRP in the media of developing ARC neurons, we used the AGRP Quantikine ELISA kit (R&D Systems, #DAGR00), which employs a solid phase sandwich ELISA that contains Sf 21-expressed recombinant human AGRP that has previously been shown to quantitate the recombinant factor. Cell culture supernatants we collected between 48-72h post media change, and they were measured in duplicates with an assay range of 7.5-500 pg/mL.

### FGF1 stimulation experiment

Human FGF1 (Miltenyi, #130-095-789) was reconstituted in water to 100 ug/ml. Accutase was pre-warmed at 37°C for 10 min. Half the media on d50 and d70 ARC cultures was removed and 100 ng/ml of FGF1 was added to the treated well for a 50ng/ml final concentration. For the control condition, the vehicle media had no FGF1. Cells received FGF1 treatment for 30 min before one wash with PBS and 150 µL of warm accutase was added and incubated at 37°C for 5 min. Cells were taken off with ice-cold NB21 medium and placed on ice before centrifugation at 700g for 10 min. Supernatant was removed and the pellet snap frozen on dry ice, kept at -80°C until snRNAseq.

### Single-cell/single-nucleus RNA sequencing sample preparation

Batch-0, batch-1, batch-2 and batch-4 d16 progenitors, which underwent scRNA sequencing, were thawed in wash media (DMEM/F12 + 5%KOSR). After counting, cells were centrifuged at 500xg for 5 min before resuspension at 5 mio cells/mL in PBS-/-with 0.5% BSA. Cells were stained with 0.5 µg of unique TotalSeq^TM^-A anti-Nuclear Hashtags (HTO) (Biolegend) and incubated for 30 min at 4°C. After antibody tagging, cells underwent three washes with PBS-/-, with final resuspension at 1000 cells/µL in PBS^-/-^ with 0.5% BSA. Samples were FACS sorted and pooled at equal ratios for one 10X lane using the 10X v3.1 chemistry kit.

The d25, 50 and 70 batch-0, batch-1, batch-2 and batch-4 were profiled using snRNA-sequencing. The samples were kept on ice and always centrifuged in cooled centrifuges. The nuclei were thawed on ice for 2 min and lysed with EZ Lysis Buffer (Nuclei EZ Prep nuclei isolation kit, Sigma, #NUC101-1KT). The lysed sample incubated on ice for 5 min, suspended in nuclei buffer (PBS-/-with 1% BSA, 2.5 mM MgCl (Sigma, #M1028), 0.2U/µL RNase inhibitor (Sigma, #3335399001)) and centrifuged for 5 min at 500xg. Then it was filtered using 40 µm cell strainer (PluriSelect, #43-10040-40) into a 2mL Protein LoBind tube (Sigma, #EP0030108132) and incubated on ice for 15 min. Next the sample was incubated together with unique TotalSeq^TM^-A anti-Nuclear Hashtags (HTO) for multiplexing for 30 min. The hashtagged nuclei from each batch were washed twice with Nuclei Buffer and stained with 7-AAD (ThermoFisher Scientific, #00-6993-50) before FACS sorting (SONY SH800S cell sorter) with a 70µm sorting chip (Sony Biotechnologies, cat. no.: LE-C3207) into a 2mL Protein LoBind tube with 18.8µL RT Reagent B (10X Genomics, Chromium Next GEM Single Cell 3’ Kit v3.1, cat. no.: PN-1000268). Following sorting, the volume was adjusted to 43.1µL with Nuclei Buffer and the final GEM Master Mix reagents were added as per manufactures procedure, which was followed from then on for library preparation with dual indexing. Each sample was divided into three 10X lanes.

HTO libraries were prepared by following the procedure from BioLegend. In short, for generating GEM cDNA, the reaction was added 1 µL of 0.2µM HTO primer (5’-GTGACTGGAGTTCAGACGTGTGC*T*C) prior to PCR cycling and cleaned up using SPRISelect magnetic beads (Beckman Coulter, #B23318), 80% ethanol, and eluted with Buffer EB (Qiagen, #1014609). The concentration was determined with Qubit (Thermo Fisher Scientific) and Qubit dsDNA HS Assay Kit (Thermo Fisher Scientific, #Q32854). Libraries reactions were generated with the following mixture: 20ng HTO cDNA, 2.5µL 10µM SI PCR primer (5’-AATGATACGGCGACCACCGAGATCTACACTCTTTCCCTACACGACGC*T*C), 2.5µL 10µM TruSeq D7XX_s (5’-CAAGCAGAAGACGGCATACGAGAT[8X]GTGACTGGAGTTCAGACGTGT*G*C), 50µL KAPA HiFi HotStart ReadyMix (Roche, #KK2602), nuclease free water to 100µL. The following PCR reaction was used: 98°C, 2 min; 15x[98°C, 20 sec; 64°C, 30 sec; 72°C, 20 sec], 72°C, 5 min. Cleanup and quantification was as above. The libraries were quantified by Qubit and TapeStation (Agilent TapeStation 4200 System) with TapeStation High Sensitivity D1000 DNA (Agilent, #5067-5584 and #5067-5585).

All sc/snRNA samples were sequenced on SP/S2 flow cells (Illumina) on NovaSeq6000. All libraries were sequenced using recommended settings aiming for a depth of 50k reads per cell (28bp R1; 10bp I1; 10bp I2; 90bp R2).

### Sequence alignment and sample assignment

The raw cDNA libraries from all single cell/nucleus samples were processed using CellRanger^3^ count pipeline v7.1.0 by aligning the day 16, 25, 50, and 70 reads against a recently published human reference with optimized genome annotations^4^. The obtained gene count matrices were corrected with CellBender^5^ v0.3.0 to remove ambient RNA. The CellRanger BAM file for d16 data was processed with STARsolo^6^ (STAR, v2.7.10b) to generate intronic and exonic counts required for RNA velocity. CITE-seq-Count^7^ v1.45 was used to identify the hashtag oligo (HTO) tags from the cell/nucleus hashing FASTQ files. To assign the sample-of-origin for each cell the HTO demultiplexing was performed using an optimized cell classification strategy (based on the HTODemux Seurat function) implemented in the COMMUNEQAID pipeline (https://github.com/CBMR-Single-Cell-Omics-Platform/COMUNEQAID).

### Single cell/nucleus data processing and cluster annotation

The cells classified as doublets or negatives and cells from incorrectly patterned batch-0 were excluded from the further analysis. The resulting count matrices were filtered and processed with Scanpy^8^ v1.9.5. Genes occurring in less than three cells, and cells with high mitochondrial content or outliers in gene counts, were removed from each batch individually by manually adjusted thresholds. To address bias arising from metabolism-related factors, ribosomal (RPS/RPL genes) and mitochondrial genes (MT-genes) were excluded from downstream analysis. The resulting data was log-normalized, and the d25 batches and the d50 and 70 batches were concatenated together. This led to three different datasets, d16, d25, and d50+70, each of which was processed and analyzed separately.

The d16 data underwent highly variable gene selection (n_top_genes=2000), dimensionality reduction (PCA and UMAP), and clustering (leidenalg v0.9.0). The d25 and d50+70 batches were further processed using Seurat^9^ v4.3.0, including highly variable gene selection, integration (RPCA), clustering, and annotation. To investigate the neuronal heterogeneity in d50-70 data, the cluster annotated as ARC neurons was subtracted and processed with Seurat as described above. The resulting clusters in all the described datasets were identified and annotated based on our previous knowledge and markers described in the literature. Clusters deemed to be transcriptionally similar were merged. Consult the available code for the exact steps and parameters used in processing, clustering, and annotation. These processed datasets (d16, d25, d50+70, d50+70 neurons) formed the basis of the rest of the analysis. Lists of differentially expressed genes (DEGs) for each cluster at each time point can be found in **Extended Data Table 1**.

### Trajectory inference with RNA velocity and Monocle3

To unravel the differentiation direction of day 16 cells, we conducted RNA velocity analysis using the stochastic model from scVelo^10^ v0.3.2. The resulting velocity stream was visualized using UMAP embedding. To further examine the lineage separation of neuronal and tanycyte lineages, we used FastMNN^11^ implemented in batchelor package v1.18.1 to integrate all d16 cells together with cells annotated as tanycytes from d25 and d50_70 datasets. Monocle3^12^ v1.3.7 was used to infer pseudotime lineages of developing tanycytes. For the identified tanycyte lineage gene expression of genes of interest was approximated along pseudotime using polynomial fit (polyfit function from numpy v1.26.4) and visualized using heatmap.

### Differential gene expression analysis

We performed differential gene expression analysis using edgeR^13^ v4.0.16. To analyze the response for FGF1 stimulation a pseudo-bulk gene expression matrix was generated by summing the transcript counts for all cells within the same cell type, differentiation, and treatment combination. Respectively, to identify genes with differential expression between cell types, we summed transcript counts within each cell type and differentiation batch. The pseudo-bulk gene expression matrix was then filtered and normalized. We used a generalized linear model implemented in glmQLFTest function (edgeR) to test for differential expression including treatment group and differentiation batch, or cell type and differentiation batch as variables in the design matrix.

### Single cell transcriptomic comparison with human data

We used a publicly available sc/snRNA-seq dataset^14^ containing samples from prenatal (gestational weeks 6 to 25) and adult human hypothalamus to unravel transcriptional similarity through reference predictions and to compare gene expression profiles observed *in vitro*. To compare gene expression profiles of canonical marker genes for ARC development and tanycyte identity we used the fetal dataset described in their study. To see whether our d50+70 dataset resembles data from human arcuate nucleus we used their fetal (neuronal and non-neuronal hypothalamic lineages) and neuronal (hypothalamic nuclei, fetal and adult) datasets. Each batch in both datasets underwent quality control, where we depleted the remaining outlier cells with automatic thresholding based on median absolute deviations (MAD) (median_abs_deviation from scipy v1.11.2) (consult the available code for MAD cutoffs). To ensure correspondence with our data, genes occurring in less than three cells, and ribosomal (RPS/RPL genes), and mitochondrial genes (MT-genes) were excluded. To integrate all batches into a human fetal and adult hypothalamic reference, we used scANVI, which was initialized with integrated scVI model from scvi-tools^15,16^ v1.1.2. scANVI projects cells into a low-dimensional space and generates a model of learned features which allows the prediction of unobserved cell types by retraining the scANVI reference model together with unseen data. Using this strategy, we applied scANVI’s functionality to predict cell types from our *in vitro* data to assess whether our data corresponds human hypothalamus. To visualize the distribution of predicted cell types, we used a bar plot to display the percentage of cells predicted to belong to certain labels. We used a Sankey plot to illustrate the correspondence between our *in vitro* cell type annotations and the predicted labels from the human reference. For the Sankey plot we only included those cell types that represented more than 1% of the total predicted cells. To project our data to the scANVI integrated reference UMAP-space we used mapscvi^17^v0.0.2 function.

### Statistical analysis

All statistical methods appoint a significance level of alpha=0.05. All quantitative values are shown as mean SEM and asterisks denotes the level of significance based on statistical test. Normality of data was assessed using Shapiro-Wilk test. Normally distributed data was analyzed with one-way ANOVA and Tukey’s multiple comparison test, if SD was unequal, Brown-Forsythe ANOVA with Dunnett’s T3 multiple comparisons test was performed. For non-normal distributions, Kruskal-Wallis test and Dunn’s multiple comparison test was used. For statistical analysis between two groups a student t-test was used, if they were non-normally distributed a Mann-Whitney test was applied.

## Data availability

All data necessary for the conclusions of the study are provided with the Article. Sc/snRNAseq data are available upon request.

## Code availability

The scripts used to analyze the RNAseq data are available at GitHub (https://github.com/kirkeby-lab/ARC_VMH_analysis).

## Extended Data

**Extended Data Table 1:** List of differentially expressed genes (DEGs) in all clusters from all time points of scRNAseq and snRNAseq datasets

## References

1. Brüning, J. C. & Fenselau, H. Integrative neurocircuits that control metabolism and food intake. Science 381, eabl7398 (2023).

2. Myers, M. G., Affinati, A. H., Richardson, N. & Schwartz, M. W. Central nervous system regulation of organismal energy and glucose homeostasis. Nat Metab 3, 737–750 (2021).

3. Saper, C. B. & Lowell, B. B. The hypothalamus. CurrentBiology 24, R1111–R1116 (2014).

4. Zhang, J., Chen, D., Sweeney, P. & Yang, Y. An excitatory ventromedial hypothalamus to paraventricular thalamus circuit that suppresses food intake. Nature Communications 11, 6326 (2020).

5. Tu, L., Fukuda, M., Tong, Q. & Xu, Y. The ventromedial hypothalamic nucleus: watchdog of whole-body glucose homeostasis. Cell Biosci 12, 71 (2022).

6. Prevot, V. et al. The Versatile Tanycyte: A Hypothalamic Integrator of Reproduction and Energy Metabolism. Endocrine Reviews 39, 333–368 (2018).

7. Bolborea, M. & Dale, N. Hypothalamic tanycytes: potential roles in the control of feeding and energy balance. Trends in Neurosciences 36, 91–100 (2013).

8. Benevento, M. et al. A brainstem–hypothalamus neuronal circuit reduces feeding upon heat exposure. Nature 628, 826–834 (2024).

9. Pan, W. W. & Myers, M. G. Leptin and the maintenance of elevated body weight. Nat Rev Neurosci 19, 95–105 (2018).

10. Dong, Y. et al. Time and metabolic state-dependent effects of GLP-1R agonists on NPY/AgRP and POMC neuronal activity in vivo. Molecular Metabolism 54, 101352 (2021).

11. Rupp, A. C. et al. Suppression of food intake by Glp1r/Lepr-coexpressing neurons prevents obesity in mouse models. Journal of Clinical Investigation 133, e157515 (2023).

12. Sisley, S. et al. Neuronal GLP1R mediates liraglutide’s anorectic but not glucose-lowering effect. J. Clin. Invest. 124, 2456–2463 (2014).

13. Ludwig, M. Q. et al. A genetic map of the mouse dorsal vagal complex and its role in obesity. Nat Metab 3, 530–545 (2021).

14. Kim, D. W. et al. The cellular and molecular landscape of hypothalamic patterning and differentiation from embryonic to late postnatal development. Nature Communications 11, 4360 (2020).

15. Kim, D. W. et al. Single-cell analysis of early chick hypothalamic development reveals that hypothalamic cells are induced from prethalamic-like progenitors. Cell Reports 38, 110251 (2022).

16. Huang, W. K. et al. Generation of hypothalamic arcuate organoids from human induced pluripotent stem cells. Cell Stem Cell 28, 1657–1670.e10 (2021).

17. Rajamani, U. et al. Super-Obese Patient-Derived iPSC Hypothalamic Neurons Exhibit Obesogenic Signatures and Hormone Responses. Cell Stem Cell 22, 698–712.e9 (2018).

18. Chen, H.-J. C. et al. Profiling human hypothalamic neurons reveals a candidate combination drug therapy for weight loss. Preprint at 10.1101/2023.07.18.549357 (2023).

19. Torz, L. et al. NPFF Decreases Activity of Human Arcuate NPY Neurons: A Study in Embryonic-Stem-Cell-Derived Model. IJMS 23, 3260 (2022).

20. Bedont, J. L., Newman, E. A. & Blackshaw, S. Patterning, specification, and differentiation in the developing hypothalamus. Wiley Interdisciplinary Reviews: Developmental Biology 4, 445–468 (2015).

21. Cobos, I., Shimamura, K., Rubenstein, J. L. R., Martínez, S. & Puelles, L. Fate Map of the Avian Anterior Forebrain at the Four-Somite Stage, Based on the Analysis of Quail–Chick Chimeras. Developmental Biology 239, 46–67 (2001).

22. Puelles, L. & Rubenstein, J. L. R. A new scenario of hypothalamic organization: Rationale of new hypotheses introduced in the updated prosomeric model. Frontiers in Neuroanatomy 9, (2015).

23. Ma, T., Wong, S. Z. H., Lee, B., Ming, G. li & Song, H. Decoding neuronal composition and ontogeny of individual hypothalamic nuclei. Neuron 109, 1150–1167.e6 (2021).

24. Padilla, S. L., Carmody, J. S. & Zeltser, L. M. Pomc-expressing progenitors give rise to antagonistic neuronal populations in hypothalamic feeding circuits. Nat Med 16, 403–405 (2010).

25. Merkle, F. T. et al. Generation of neuropeptidergic hypothalamic neurons from human pluripotent stem cells. Development (Cambridge) 142, 633–643 (2015).

26. Sarrafha, L. et al. Novel human pluripotent stem cell-derived hypothalamus organoids demonstrate cellular diversity. iScience 26, 107525 (2023).

27. Chinnaiya, K. et al. A neuroepithelial wave of BMP signalling drives anteroposterior specification of the tuberal hypothalamus. eLife 12, e83133 (2023).

28. Herb, B. R. et al. Single-cell genomics reveals region-specific developmental trajectories underlying neuronal diversity in the human hypothalamus. Science Advances 9, eadf6251.

29. Steuernagel, L. et al. HypoMap—a unified single-cell gene expression atlas of the murine hypothalamus. Nat Metab 4, 1402–1419 (2022).

30. Jais, A. et al. PNOCARC Neurons Promote Hyperphagia and Obesity upon High-Fat-Diet Feeding. Neuron 106, 1009–1025.e10 (2020).

31. Huisman, C. et al. Single cell transcriptome analysis of developing arcuate nucleus neurons uncovers their key developmental regulators. Nature Communications 10, (2019).

32. Scarlett, J. M. et al. Central injection of fibroblast growth factor 1 induces sustained remission of diabetic hyperglycemia in rodents. Nature Medicine 22, 800–806 (2016).

33. Bentsen, M. A. et al. Transcriptomic analysis links diverse hypothalamic cell types to fibroblast growth factor 1-induced sustained diabetes remission. Nat Commun 11, 4458 (2020).

34. Alvarez-Bolado, G., Paul, F. A. & Blaess, S. Sonic hedgehog lineage in the mouse hypothalamus: from progenitor domains to hypothalamic regions. Neural Dev 7, 4 (2012).

35. Skidmore, J. M., Cramer, J. D., Martin, J. F. & Martin, D. M. Cre fate mapping reveals lineage specific defects in neuronal migration with loss of Pitx2 function in the developing mouse hypothalamus and subthalamic nucleus. Molecular and Cellular Neuroscience 37, 696–707 (2008).

36. López-González, L., Martínez-de-la-Torre, M. & Puelles, L. Populational heterogeneity and partial migratory origin of the ventromedial hypothalamic nucleus: genoarchitectonic analysis in the mouse. Brain Struct Funct 228, 537–576 (2023).

37. Kim, D. W. et al. The cellular and molecular landscape of hypothalamic patterning and differentiation from embryonic to late postnatal development. Nature Communications 11, (2020).

38. Caron, A. et al. POMC neurons expressing leptin receptors coordinate metabolic responses to fasting via suppression of leptin levels. eLife 7, e33710 (2018).

39. Biglari, N. et al. Functionally distinct POMC-expressing neuron subpopulations in hypothalamus revealed by intersectional targeting. Nature Neuroscience 24, 913–929 (2021).

40. Yao, Z. et al. A Single-Cell Roadmap of Lineage Bifurcation in Human ESC Models of Embryonic Brain Development. Cell Stem Cell 20, 120–134 (2017).

41. Sotelo-Hitschfeld, T. et al. GABAergic disinhibition from the BNST to PNOC ARC neurons promotes HFD-induced hyperphagia. Cell Reports 43, 114343 (2024).

42. Lehr, S. et al. Self-organised pattern formation in the developing neural tube by a temporal relay of BMP signalling. Preprint at 10.1101/2023.11.15.567070 (2023).

43. Manning, L. et al. Regional Morphogenesis in the Hypothalamus: A BMP-Tbx2 Pathway Coordinates Fate and Proliferation through Shh Downregulation. Developmental Cell 11, 873–885 (2006).

44. Campbell, J. N. et al. A molecular census of arcuate hypothalamus and median eminence cell types. Nature Neuroscience 20, 484–496 (2017).

45. Brown, J. M. et al. Role of hypothalamic MAPK/ERK signaling and central action of FGF1 in diabetes remission. iScience 24, 102944 (2021).

## Methods references

1. Pantazis, C. B. et al. A reference human induced pluripotent stem cell line for large-scale collaborative studies. Cell Stem Cell 29, 1685–1702.e22 (2022).

2. Nolbrant, S., Heuer, A., Parmar, M. & Kirkeby, A. Generation of high-purity human ventral midbrain dopaminergic progenitors for in vitro maturation and intracerebral transplantation. Nat Protoc 12, 1962–1979 (2017).

3. Zheng, G. X. Y. et al. Massively parallel digital transcriptional profiling of single cells. Nat Commun 8, 14049 (2017).

4. Pool, A.-H., Poldsam, H., Chen, S., Thomson, M. & Oka, Y. Recovery of missing single-cell RNA-sequencing data with optimized transcriptomic references. Nat Methods 20, 1506–1515 (2023).

5. Fleming, S. J. et al. Unsupervised removal of systematic background noise from droplet-based single-cell experiments using CellBender. Nat Methods 20, 1323–1335 (2023).

6. Kaminow, B., Yunusov, D. & Dobin, A. STARsolo: accurate, fast and versatile mapping/quantification of single-cell and single-nucleus RNA-seq data. Preprint at 10.1101/2021.05.05.442755 (2021).

7. Stoeckius, M. et al. Cell Hashing with barcoded antibodies enables multiplexing and doublet detection for single cell genomics. Genome Biol 19, 224 (2018).

8. Wolf, F. A., Angerer, P. & Theis, F. J. SCANPY: large-scale single-cell gene expression data analysis. Genome Biol 19, 15 (2018).

9. Hao, Y. et al. Integrated analysis of multimodal single-cell data. Cell 184, 3573–3587.e29 (2021).

10. Bergen, V., Lange, M., Peidli, S., Wolf, F. A. & Theis, F. J. Generalizing RNA velocity to transient cell states through dynamical modeling. Nat Biotechnol 38, 1408–1414 (2020).

11. Haghverdi, L., Lun, A. T. L., Morgan, M. D. & Marioni, J. C. Batch effects in single-cell RNA-sequencing data are corrected by matching mutual nearest neighbors. Nat Biotechnol 36, 421–427 (2018).

12. Cao, J. et al. The single-cell transcriptional landscape of mammalian organogenesis.Nature 566, 496–502 (2019).

13. Chen, Y., Chen, L., Lun, A. T. L., Baldoni, P. L. & Smyth, G. K. edgeR 4.0: powerful differential analysis of sequencing data with expanded functionality and improved support for small counts and larger datasets. Preprint at 10.1101/2024.01.21.576131 (2024).

14. Herb, B. R. et al. Single-cell genomics reveals region-specific developmental trajectories underlying neuronal diversity in the human hypothalamus. Science Advances 9, eadf6251.

15. Gayoso, A. et al. A Python library for probabilistic analysis of single-cell omics data. Nat Biotechnol 40, 163–166 (2022).

16. Xu, C. et al. Probabilistic harmonization and annotation of single-cell transcriptomics data with deep generative models. Molecular Systems Biology 17, e9620 (2021).

17. Steuernagel, L. et al. HypoMap—a unified single-cell gene expression atlas of the murine hypothalamus. Nat Metab 4, 1402–1419 (2022).

