## Supplementary tables for "Generation of human appetite-regulating neurons and tanycytes from stem cells"

[1]Novo Nordisk Foundation Center for Stem Cell Medicine (reNEW), Faculty of Health and Medical Sciences, University of Copenhagen, 2200, Copenhagen, Denmark
[2]Department of Experimental Medical Sciences, Wallenberg Centre for Molecular Medicine (WCMM) and Lund Stem Cell Centre, Lund University- SE-221 84 Lund, Sweden
[3]Department of Neuroscience, Faculty of Health and Medical Sciences, University of Copenhagen, 2200, Copenhagen, Denmark
[4]Novo Nordisk Foundation Center for Basic Metabolic Research (CBMR), Faculty of Health and Medical Sciences, University of Copenhagen, 2200, Copenhagen, Denmark
[*] These authors contributed equally

### Supplementary information

| Gene symbol | Gene name | Forward primer sequence | Reverse primer sequence |
|---|---|---|---|
| *ACTB* | Actin beta | ATGTGGCCGAGGACTTTGATTG | ATGGCAAGGGACTTCCTGTAAC |
| *AGRP* | Agouti related neuropeptide | TCCCTGTCCTGTGGAAATTTGT | TCTCCAATTTGGGGTGAGGTTT |
| *AQP4* | Aquaporin 4 | TTGGACCTGCAGTTATCATGGG | CTATGATGGGCCCAACCCAATA |
| *ARX* | Aristaless related homeobox | CCTGAGCACTTTCCTCGGAGCG | TGGAAAAGAGCCTGCCGAATGCC |
| *BRN2* | POU3F2, POU class 3 homeobox 2 | ATGCGCGGCTCCTTTAACCGG | TTAGACGCTGCGGTCGCCATG |
| *BSX* | Brain specific homeobox | GAGAAGAGGTTCGAGATCCAGC | GTTCTGGAACCACGTTTTCACC |
| *CART* | CART prepropeptide | GCGTCCATTCTCCTCCATACAT | GTTGCTTAAGCCAAACTCCAGG |
| *CRH* | Corticotropin releasing hormone | CTGTACCATAGCGCTGCTCTTA | TTGTGCATGCTAAGTAAGGGGT |
| *CITED1* | Cbp/p300 interacting transactivator with Glu/Asp rich carboxy-terminal domain 1 | AGTGGATGAGGAAGTGCTGATG | AAGTCAAACTCATTCTGCCCCA |
| *COL1A1* | Collagen type I alpha 1 chain | CTTATGAAACCCCAATGCTGCC | GGGAGACAGATTTGGGAAGGAG |
| *CRYM* | Crystallin mu | ACAGAGCCCATTTTGTTTGGTG | CATCCAGTTCTCTCCAGTCAGG |
| *DBX1* | Developing brain homeobox 1 | GAAGTTTGGAGTGAACGCCATC | GACCCTTCGAAGTAGGGAAAGG |
| *DIO2* | Iodothyronine Deiodinase 2 | GTTATAAGGCAACCCCCGGTAT | AACCTCAGCTAATGGGACCAAG |
| *DLX2* | Distal-less homeobox 2 | ACCAGACCTCGGGATCCGCC | CTGCGGGGTCTGAGTGGGGT |
| *EN1* | Engrailed homeobox 1 | CGTGGCTTACTCCCCATTTA | TCTCGCTGTCTCTCCCTCTC |
| *FEZF1* | FEZ family zinc finger 1 | GGTACATTCCACATTCGTGAGC | TCACGTGCAATAATCAAAACCA |
| *FGF10* | Fibroblast growth factor 10 | GTTGCTGTTCTTGGTGTCTTCC | TGACACCATGTCCTGACCAAG |
| *FOXA1* | Forkhead box A1 | GGGCAGGGTGGCTCCAGGAT | TGCTGACCGGGACGGAGGAG |
| *FOXA2* | Forkhead box A2 | CCGTTCTCCATCAACAACCT | GGGGTAGTGCATCACCTGTT |
| *FOXB1* | Forkhead box B1 | GTGGTCGGACTTAAGCACCTT | GTGGTCGGACTTAAGCACCTT |
| *FOXD1* | Forkhead box D1 | CCTGTCCAGTGTCGAGAACTTT | AACCACCAAGACGAGAAAAGGA |
| *FOXG1* | Forkhead box G1 | TCAACGGCATCTACGAGTTCAT | AAGCACTTGTTGAGGGACAGAT |
| *GAPDH* | Glyceraldehyde-3-phosphate dehydrogenase | TTGAGGTCAATGAAGGGGTC | GAAGGTGAAGGTCGGAGTCA |

| | | | |
|---|---|---|---|
| *GBX2* | Gastrulation brain homeobox 2 | GTTCCCGCCGTCGCTGATGAT | GCCGGTGTAGACGAAATGGCCG |
| *GHRH* | Growth hormone releasing hormone | AGGAACTCCCAGGGATGAAGAT | CAGTTGCATTTTGGCTACAGGT |
| *GRH* | Growth hormone receptor | ACTAGCAATGGTGGTACAGTGG | TCAGTAAAGTCCAGTTGAGGGC |
| *GPR149* | G protein-coupled receptor 149 | GACTGGGAGTGGTGTAGGAGTA | GGCATAACCGGAACGCTGAC |
| *HCRT* | Hypocretin neuropeptide predursor | CCTCAAGGTTCCTGGCTTTTTG | GGAAGGAAGGTTCATGGTGTCT |
| *ISL1* | ISL LIM homeobox 1 | AAGCGCAGGAAGAGAGACTG | CCAAGAGACCCAGGATTTCA |
| *KISS1* | KiSS-1 metastasis suppressor | TGAACTTCAGACCCCAAAGGAG | TCTTTTATTGCCTCGGGTTGGA |
| *LEPR* | Leprin receptor | TACTGTTACGGTTCTGGCCATC | TGCTCATAGGCCATGAAAAGGT |
| *LHX2* | LIM homeobox 2 | GGGCGACCACTTCGGCATGAA | CGTCGGCATGGTTGAAGTGTGC |
| *LHX6* | LIM Homeobox 6 | AGGCAAGAACATCTGCTCCAG | GCCAGATGAGGTTGTTGACCTT |
| *LHX8* | LIM Homeobox 6 | AGGCAAGAACATCTGCTCCAG | GCCAGATGAGGTTGTTGACCTT |
| *LHX9* | LIM Homeobox 9 | TGCCTGAAGTGCTGTGAATGTA | TTGCAGTAAATGCTACCGTCCT |
| *MASH1* | ASCL1, achaete-scute family bHLH transcription factor 1 | CTAAAGATGCAGGTTGTGCG | GGAGCTTCTCGACTTCACCA |
| *MC4R* | Melancortin 4 receptor | CTTTTTCATCTGCAGCTTGGCT | TGTGAAACTCTGTGCATCCGTA |
| *MCH* | Melanin concentrating hormone | TACATTCAGGTTGGGGAAAGGC | CCAGGGAAGGAGCAATAACTGA |
| *MPZ* | Myelin protein zero | CTCAGGTCACGCTGTATGTCTT | GAACCACGTAGAAAAGCAGCAG |
| *NCAM1* | Neural cell adhesion molecule 1 | GTCAGAGGCCACCGTCAACGTG | CTTCCCCCTCCCGGAACTCCTG |
| *NGN3* | Neurogenin 3 | CTGAACTTGGCGACCAGAAGC | TTGAGGCGTCATCCTTTCTACC |
| *NHLH2* | Nescient helix-loop-helix 2 | ACCCACTGGAGACTTTGAGTTC | GCATACTCTGAACTTCTGCCCT |
| *NKX2.1* | NK2 homeobox 1 | AGGGCGGGGCACAGATTGGA | GCTGGCAGAGTGTGCCCAGA |
| *NPY* | Neuropeptide Y | GAAAATGTTCCCAGAACTCGGC | TAGGAAAAGGCCAGAGAGCAAG |
| *NR5A1* | Nuclear receptor subfamily 5 group A member 1 | GGAACAAGTTTGGGCCGATGTA | GTGCCTTCTTCTGCTGTTTCAG |
| *NROB1* | Nuclear receptor subfamily 0 group B member 1 | GCCATCAAGTGCTTTCTTTCCA | TAGGCGTACTCCTTGGTACTGA |
| *NTS* | Homo sapiens neurotensin | GCTCAAAGTACTACAGCAAAGCC | GGCGCTATTACTTTGTTTTGGGT |
| *ONECUT2* | One cut homeobox 2 | AAGGGGTAGAGCTGGTGTATCT | TGTTGTTTCAGGGGTGACTTGA |
| *OTP* | Orthopedia homeobox | TAGAAGGGAAGGCTTCTCAGGA | GTCAGATCACCTCTTCCTCGTC |

| | | | |
|---|---|---|---|
| *OTX2* | Orthodenticle homeobox 2 | ACAAGTGGCCAAT TCACTCC | GAGGTGGACAAGGGA TCTGA |
| *PAX6* | Paired box 6 | TGGTATTCTCTCC CCCTCCT | TAAGGATGTTGAACGG GCAG |
| *PCSK1* | Proprotein convertase subtilisin/kexin type 1 | GCAATGCCCGTAA TGCTTAGAG | TTCCAAGGACAGAGT GATTCCG |
| *PCSK2* | Proprotein convertase subtilisin/kexin type 2 | GATTGACTATCTCC ACCCGGAC | TCTGTGTACCGAGGG TAAGGAT |
| *PDYN* | Prodynorphin | CTCATTCCCAGGC ACTCTCTTT | TTTCCTCTCCTATCCA GCCTCA |
| *PITX2* | Paired like homeodomain 2 | AACTCTATGAACG TCAACCCCC | CGACATGCTCATGGA CGAGATA |
| *POMC* | Proopiomelanocortin | TTTCATGACCTCC GAGAAGAGC | GATGATGGCGTTTTTG AACAGC |
| *PRDM12* | PR/SET domain 12 | GTGGGAGGTGTTC AATGAGGAT | GTTCGTTACGTGCACA CTTGAT |
| *PVALB* | Parvalbumin | GGACAAGGACAAA AGTGGCTTC | CAGCCATCAGCATCTT GGTTTC |
| *RAX* | Retina and anterior neural fold homebox | CCTCTCAGTTCAC CAAGCAGAT | TGATCAACCTTGGGT GTTAGGG |
| *RFX4* | Regulatory factor X4 | ACCTTGCCATCTG TCTTGTCAT | ATAGGGATGGTACCAC CAGGAA |
| *SHH* | Sonic hedgehog | CCAATTACAACCC CGACATC | AGTTTCACTCCTGGC CACTG |
| *SIM1* | Single-minded family bHLH transcription factor 1 | AAAGGGGGCCAA ATCCCGGC | TCCGCCCCACTGGCT GTCAT |
| *SIM2* | SIM bHLH transcription factor 2 | GGCTACTTGAAGA TCAGGCAGT | ATCTGGTAGCAGGAG TCGTACA |
| *SIX3* | SIX homeobox 3 | ACCGGCCTCACTC CCACACA | CGCTCGGTCCAATGG CCTGG |
| *SIX6* | SIX homeobox 6 | CTCAACAAGAATG AGTCGGTGC | ACTCCTTGGTGAACTT GTGGTT |
| *SOX14* | SRY-box 14 | CATACATCGATGAA GCCAAGCG | CTGTCCTTCTTGAGCA GGTTCT |
| *SP8* | Sp8 transcription factor | CCTGTCTGTCCGG ACTTCAA | AGGGGCAGAAACAGA AAGAGAC |
| *SP9* | Sp9 transcription factor | TCTGGCCCCAACG ACTCTTAG | CTCGTTCGTTCTCGGT GTCTC |
| *SST* | Somatostatin | GGAACCTGAAGAT CTGTCCCAG | ATAGCCGGGTTTGAGT TAGCAG |
| *TBX3* | T-box transcription factor 3 | GGGGGTAGGAGTT CCAACATTT | GCACTGAGGGAGATG TCTTTGA |
| *TH* | Tyrosine hydroxylase | CGGGCTTCTCGGA CCAGGTGTA | CTCCTCGGCGGTGTA CTCCACA |
| *TRH* | Thyrotropin releasing hormone | TCCTGGATGACCT GAGTAGGAG | TCAGGGAAAACTGGG TTCACTC |
| *VSX2* | Unc-5 netrin receptor D | CCTCTGCCCTCTG TAAATGTGT | AGAGACCTCTGCGAG AACTTTG |

**Supplementary Table 1:** List of qRT-PCR primers.

| Antigen | Host species | Dilution | Manufacturer (#Cat) |
|---|---|---|---|
| NKX2-1 | Rabbit | 1:100-200 | Abcam (#ab1333737) |
| PAX6 | Mouse | 1:1000 | Sigma-Merck (#AMAb91372) |
| AGRP | Rabbit | 1:1000 | Phoenix Pharma (#H-003-53) – discontinued |
| AGRP | Goat | 1:100-200 | R&D Systems AF634 |
| POMC (aMSH) | Sheep | 1:1000 | Millipore (#AB5087) |
| MAP2 | Mouse | 1:1000 | Sigma (#M1406) |
| TRH | Rabbit | 1:1000 | Thermo Fisher (PA5-57331) |
| TH | Mouse | 1:1000 | Immunostar (173-22941) |
| GHRH | Rabbit | 1:200 | Abcam (ab18751 |
| OTP | Rabbit | 1:500 | GeneTex (GTX119601) |
| FOXG1 | Rabbit | 1:500 | Abcam (#ab18259) |
| CRH | Rabbit | 1:1000 | Proteintech (10944-1-AP) |
| AQP4 | Rabbit | 1:1000 | Sigma-Merck (#HPA014784) |
| hNCAM1 | Mouse | 1:1000 | Santa Cruz Biotechnology #SC-106 |
| HuNu | Mouse | IF: 1:1000<br>ISH: 1:200 | Millipore #AB1281 |
| NPY | Sheep | 1:500 | Millipore #AB1583 |
| S100b | Mouse | 1:1000 | Sigma (#S2532) |
| Somatostatin | Mouse coupled-Alexa Fluor 488 | 1:500 | BD Biosciences (#566032) |
| NFIA | Rabbit | 1:200 | Abcam (#ab228897) |
| Vimentin | Chicken | 1:500 | Millipore (#AB5733) |
| NR5A1 | Mouse | 1:1000 | Thermo Fisher (#434200) |

**Supplementary Table 2:** List of primary antibodies.

| ANTIGEN | FLUOROPHORE | DILUTION | MANUFACTURER (#CAT) |
|---|---|---|---|
| RABBIT | Alexa488 | 1:200-500 | Jackson Imm. Res. (#711-545-152) |
| MOUSE | Alexa647 | 1:200-500 | Jackson Imm. Res. (#711-605-151) |
| SHEEP | Alexa Cy3 | 1:200-500 | Jackson Imm. Res. (#713-165-147) |
| MOUSE | Alexa Cy3 | 1:200 | Jackson Imm. Res. (#715-165-151) |
| MOUSE | Alexa 488 | 1:200-500 | Jackson Imm. Res. (#715-545-150) |
| RABBIT | Alexa CY3 | 1:200 | Jackson Imm. Res. (#711-165-152) |
| GOAT | Alexa488 | 1:200-500 | Jackson Imm. Res. (#705-545-147) |
| SHEEP | Alexa647 | 1:200-500 | Jackson Imm. Res. (#713-605-147) |
| GOAT | Alexa647 | 1:200 | Jackson Imm. Res. (#705-605-147) |
| MOUSE | Biotinylated | 1:500 | Vector Laboratories #BA9200 |
| DAPI | - | 1:500 | Thermo (#D3571) |

**Supplementary table 3.** List of secondary antibodies.

| Probe | Host species | Dilution | Manufacturer (#Cat) |
|---|---|---|---|
| Hs-RAX-C1 | Homo sapiens | Ready-to-use | ACD Bio-Techne (#579951) |
| Hs-TBX3-C2 | Homo sapiens | 1:50 | ACD Bio-Techne (#557441) |
| Hs-SIM1-C3 | Homo sapiens | 1:50 | ACD Bio-Techne (#544281) |
| Hs-POU3F2-C2 | Homo sapiens | 1:50 | ACD Bio-Techne (#543011) |
| Hs-PNOC-C2 | Homo sapiens | 1:50 | ACD Bio-Techne (#1045241) |
| **Fluorophore** | | | |
| Opal *480* | | 1:500 | Akoya Bio FP1500001KT |
| Opal *570* | | 1:500 | Akoya Bio FP1488001KT |
| Opal *690* | | 1:500 | Akoya Bio FP1497001KT |

**Supplementary table 4.** List of RNAscope probes and fluorophores.
