## Supplementary figures for "Generation of human appetite-regulating neurons and tanycytes from stem cells"

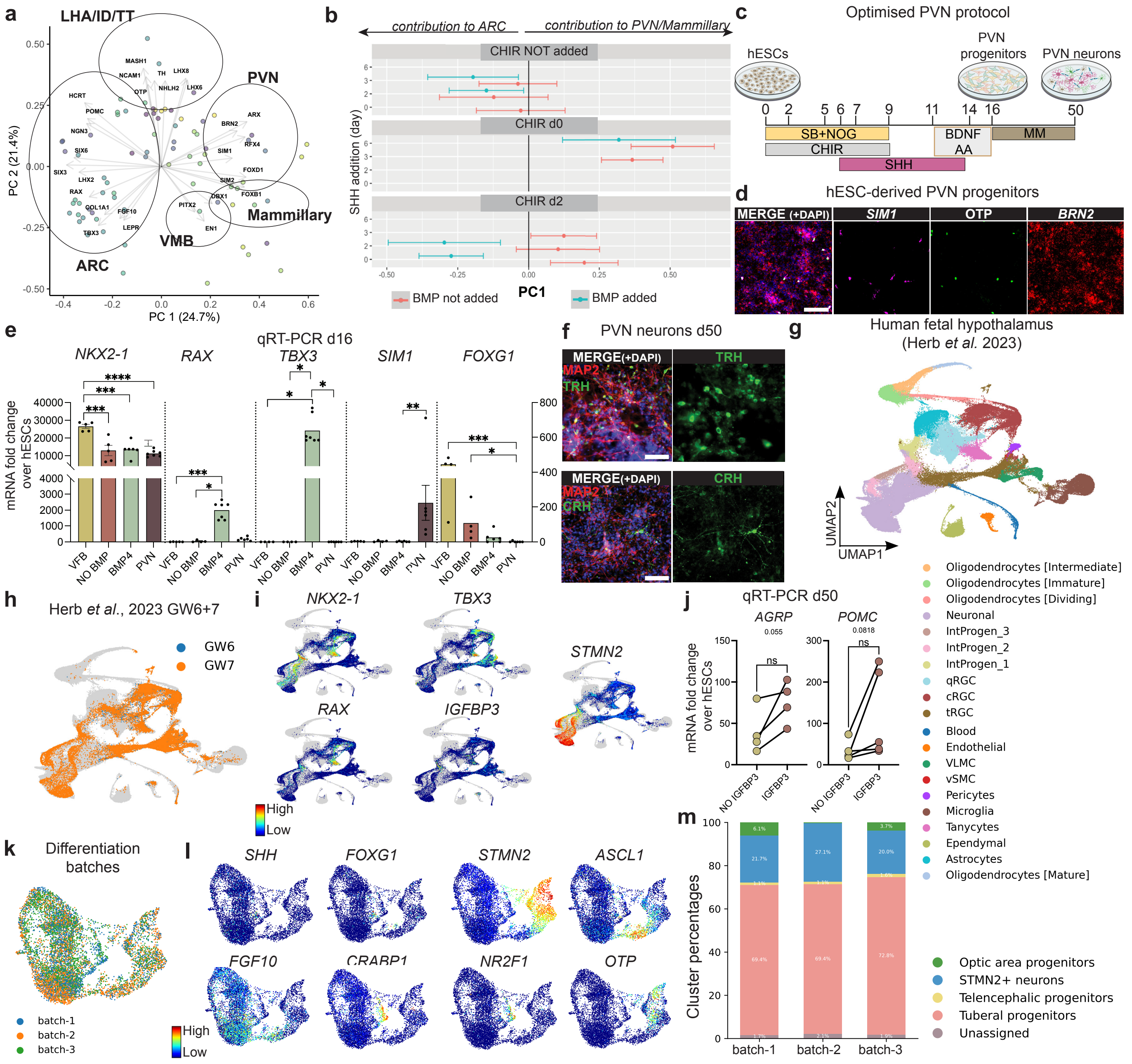


**Supplementary Fig. 1.**

**a**, PCA analysis of d16 qRT-PCR data from differentiations (n=113) with various different morphogens showed clustering of samples indicative of different regions of the hypothalamus. Each dot represents one differentiation. **b**, Estimated marginal means (EMM) to identify the average effects of addition of SHH, WNTa (CHIR), and BMP to patterning of differentiation, adjusted for other covariates in the model. The plot shows the contribution of each growth factor (added at different days) to PC1. A low PC1 score correlates with ARC-related gene expression and high PC1 score correlates PVN-related expression. **c**, qRT-PCR expression data of d16 PVN and ARC differentiations with and without BMP, including a ventral forebrain (vFB) differentiation as control. One-way ANOVA with Tukey's multiple comparisons test or Kruskal-Wallis with Dunn's multiple comparisons test. NKX2-1: VFBvsNO BMP p= 0.0003, VFBvsBMP p= 0.0003, VFBvsPVN p= <0.0001. RAX: VFBvsBMP p= 0.0006, NOBMPvsBMP p= 0.0156. TBX3: VFBvsBMP p= 0.0280, NOBMPvsBMP p= 0.0341, BMPvsPVN p= 0.0225. SIM1: BMP4vsPVN p= 0.0079. FOXG1: VFBvsPVN p= 0.0010, NO BMPvsPVN p= 0.0466. **d**, Schematic overview of the optimised PVN protocol. **e**, Combinatorial immunocytochemistry (ICC) (OTP) and *in situ* hybridization (ISH) (*SIM1* and *BRN2*) on day (d) 16 PVN progenitors. Scale bar, 100 µm. **f**, ICC analysis of d50 PVN cultures identifying the expression of MAP2, TRH and CRH. Scale bar, 100 µm. **g**, UMAP with annotations of integrated human fetal data [22]. **h**, UMAP of scRNAseq data from gestasional week (GW) 6 and 7[22]. **i**, Feature plots for key genes from the GW6 and 7 dataset showing IGFBP3 expression pattern[22]. **k**, UMAP clustering of the three ARC batches (batch 1, batch 2, batch 3) analysed at d16. **l**, Feature plots of relevant markers of d16 dataset. **j**, qRT-PCR data of d50 ARC cultures with or without IGFBP3 addition. Pairwise analysis within the same differentiation experiment, with and without treatment. Paired t-test: AGRP p= 0.055. POMC p=0.0818. **m**, Cluster percentages of selected clusters of each of the three batches. AA, ascorbic acid; ARC, arcuate nucleus; BDNF, bone-derived neurotrophic factor; MM, maturation medium; NOG, Noggin; PVN, paraventricular nucleus; SB, SB-431542; SHH, Sonic hedgehog; ID/TT, intrahypothalamic diagonal/tuberomamillary terminal; LHA, lateral hypothalamic area; VMB, ventral midbrain.

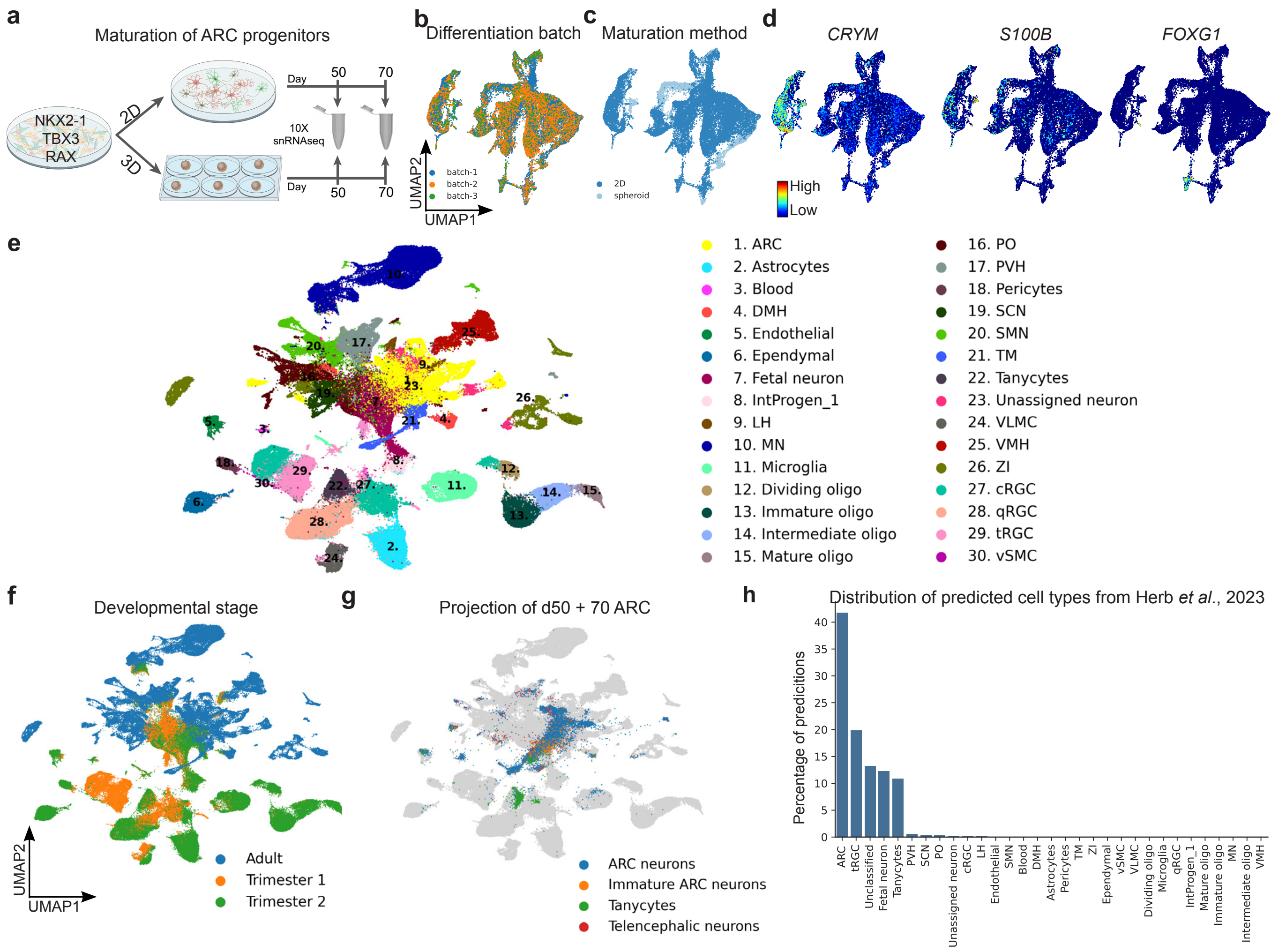


**Supplementary Fig. 2: a**, Schematic overview of 2D and 3D *in vitro* maturation of ARC progenitors. Nuclei were extracted for snRNAseq at day (d) 50 + d70. **b**, UMAP of d50 + 70 ARC neuronal data with the three batches annotated. **c**, UMAP of d50 + 70 ARC dataset showing the contribution of cells from 2D versus 3D culture. **d**, Feature plots for additional key markers of tanycytes (*CRYM* and *S100b*) and the telencephalon (*FOXG1*). **e**, UMAP of human fetal dataset with annotated clusters[28]. **f**, UMAP of the Herb *et al.* 2023[28] dataset showing contributions of cells from developmental stages. **g**, Projections of in-house d50 + 70 ARC dataset onto the Herb *et al.* 2023[28] reference. **h**, Bar plot showing the percentage of predictions d50 + 70 ARC data points to the Herb *et al.* 2023[28] reference covering all hypothalamic clusters.

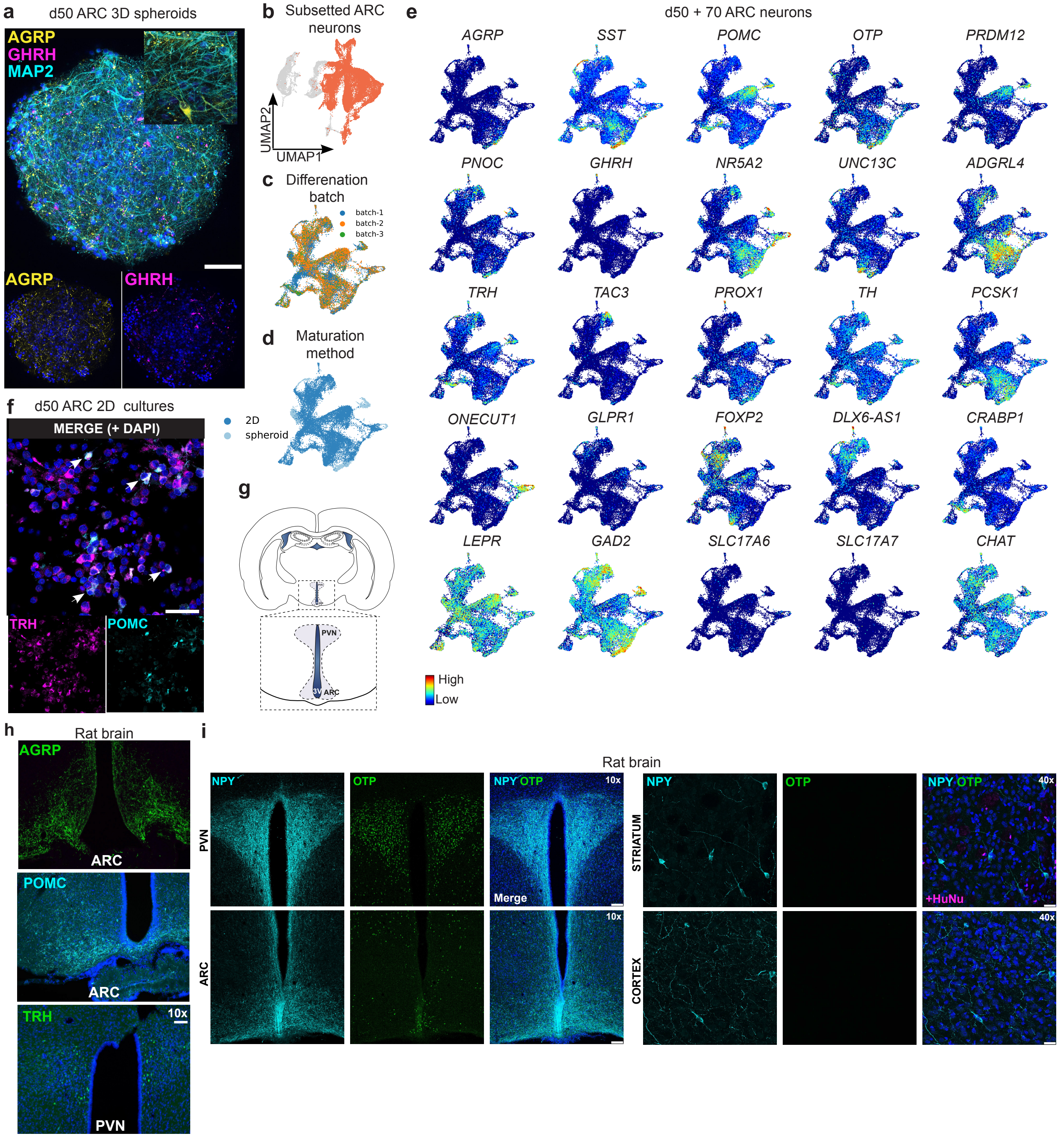


**Supplementary Fig. 3.**
**a,** Immunocytochemistry (ICC) images of d50 3D spheroid cultures depicting AGRP and GHRH. Scale bar, 100 µm. **b**, UMAP of subsetted arcuate nucleus (ARC) neurons from d50+70 late stage data for subclustering. **c**, Contributions of all three batches to subclustered ARC neurons. **d**. Contributions of 2D versus 3D maturation method to the ARC neuron subclustered dataset. **e**, Feature plots of genes of interest. **f**, ICC images of d50 2D cultures showing co-staining of TRH and POMC (αMSH) neurons. Scale bar, 50 µm. **g**, Schematic of a coronal brain section showing the location of the ARC and the paraventricular nucleus (PVN) in the rat brain. **h**, AGRP, POMC, and TRH antibody staining on rat hypothalamic brain sections as positive control for antibody specificity. Scale bars, 100 µM. **i**, NPY and OTP antibody staining of rat brain sections showing antibody specificity including a distinctly different subcellular NPY localisation in ARC NPY neurons versus striatal and cortical NPY interneurons. Scale bars, 100 µM (ARC, PVN) and 25 µM (striatum, cortex).

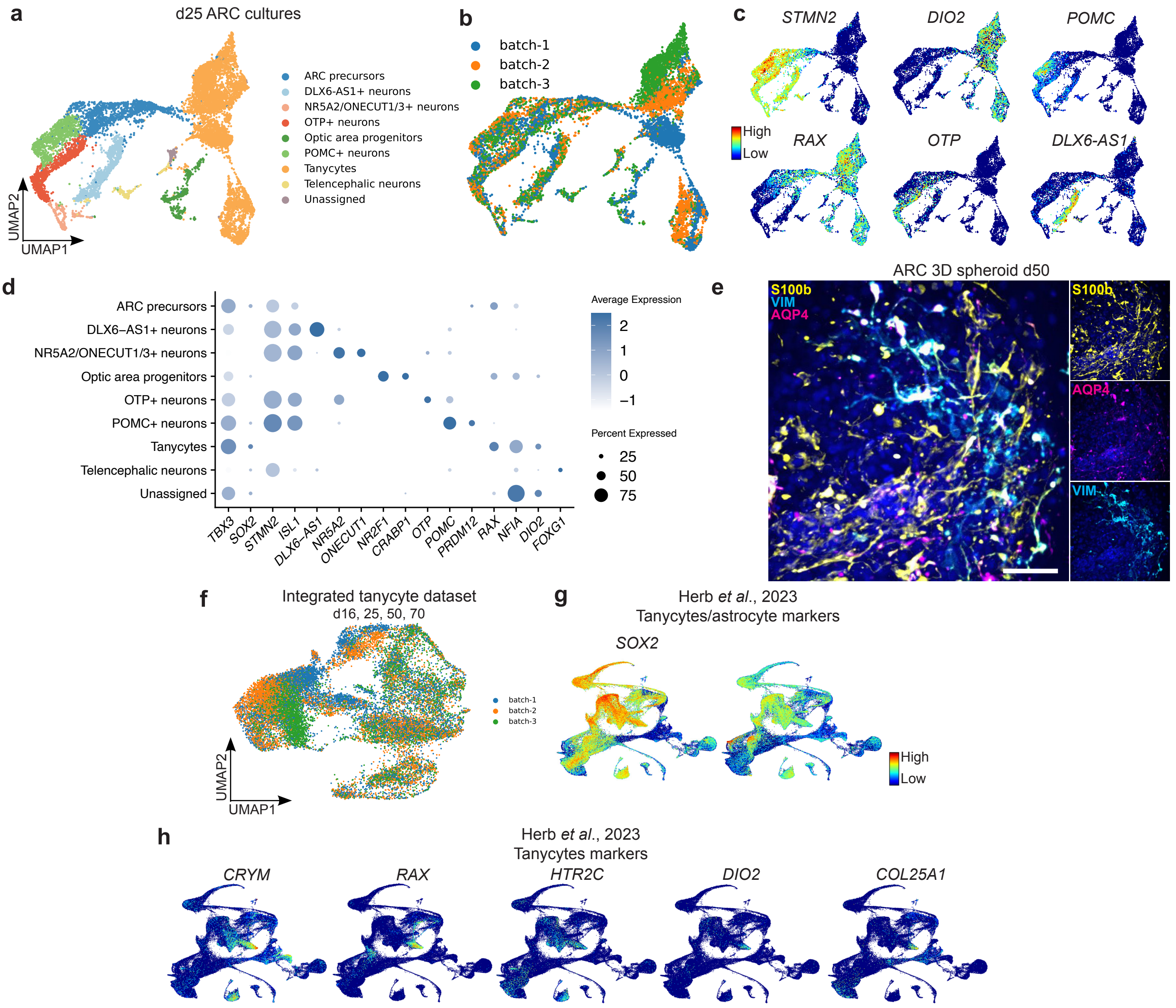


**Supplementary Fig. 4.**

**a**, UMAP of snRNAseq data of the arcuate nucleus (ARC) protocol (n=3 batches) analysed at day (d) 25. **b**, UMAP of contribution of each ARC batch to the d25 dataset. **c**, Feature plots for relevant markers in the d25 ARC dataset. **d**, Dot plot of key genes for annotated clusters of the d25 ARC dataset. **e**, ICC of d50 3D culture showing expression of AQP4, S100b, and vimentin (VIM) (dashed square). Scale bar: 50 µm. **f**, UMAP of the integrated tanycyte dataset ( d16 dataset together with the tanycyte clusters from d25, 50, and 70), showing the contribution from each batch. **g-h**, Feature maps of genes found in the stem cell-derived tanycytes from the Herb *et al*. 2023[22] reference showing several genes with selectivity for the tanycyte population in the fetal dataset (see annotations in Supplementary Fig. 1g).

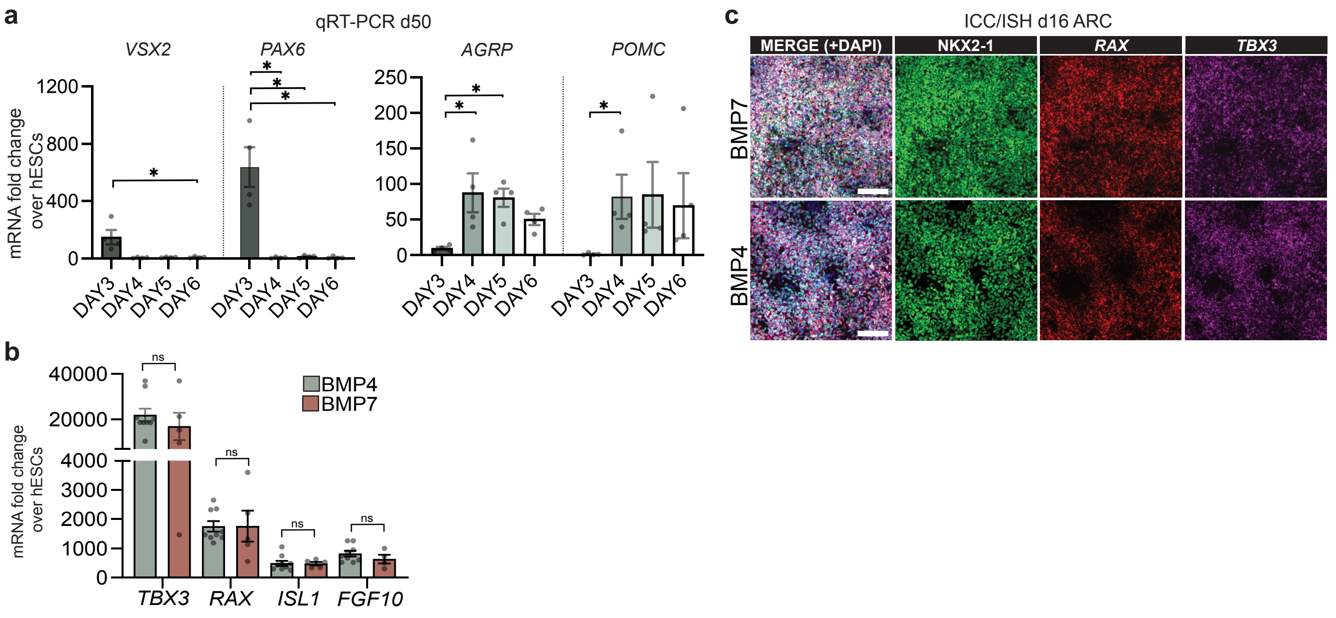


**Supplementary Fig. 5.**
**a**, qRT-PCR expression data of d50 arcuate nucleus (ARC) cultures with different BMP addition start timepoints. One-way ANOVA with Tukey's multiple comparisons test: *AGRP*: DAY3vsDAY4 *p= 0.0190, DAY3vsDAY5 *p= 0.0332. Brown-Forsythe ANOVA with Dunnett's T3 multiple comparisons test: *PAX6*: DAY3vsDAY4 *p=0.0190, DAY3vsDAY5 *p=0.0191, DAY3vsDAY6 *p=0.0191. Kruskal-Wallis with Dunn's multiple comparisons test: *VSX2*: DAY3vsDAY6 *p= 0.0360; *POMC*: DAY3vsDAY4 *p= 0.0227. **b**, qRT-PCR analysis of d16 ARC progenitors with either BMP4 or BMP7 treatment. Unpaired t-test: ns, non significant. **c**, Combinatorial immunocytochemistry (ICC) (NKX2-1) and *in situ* hybridisation (ISH) (*RAX* and *TBX3*) on d16 ARC cultures comparing BMP4 and BMP7 treatments. Scale bars, 100 μm. BMP, bone morphogenetic protein.

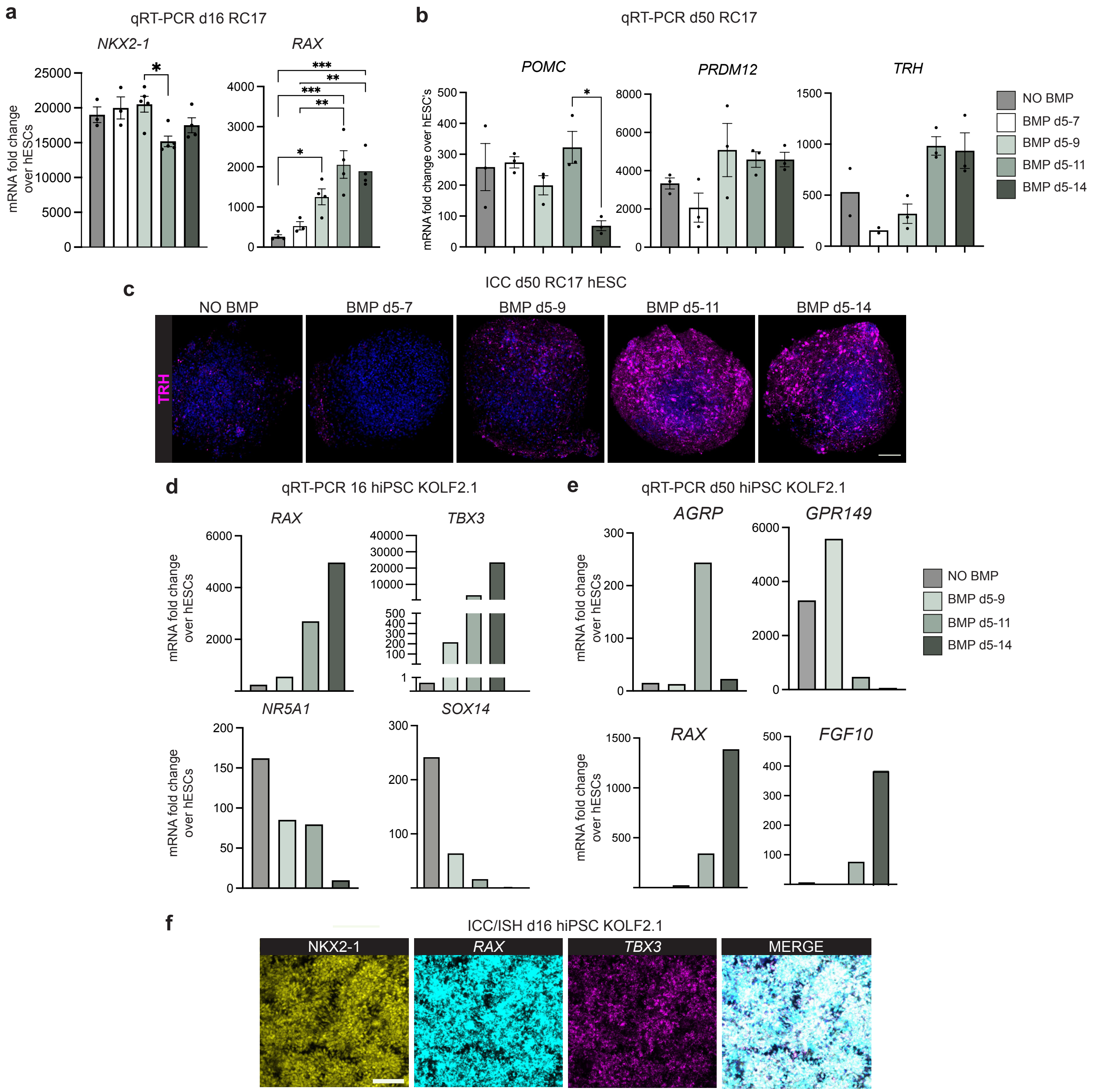


**Supplementary Fig. 6**

**a+b**, qRT-PCR of day (d) 16 (**a**) and d50 (**b**) arcuate nucleus (ARC) progenitors differentiated from RC17 hESCs with different termination days of BMP4 treatment. Kruskal-Wallis with Dunn's multiple comparisons test: *NKX2-1*: d5-9vsd5-11 *p= 0.0328. One-way ANOVA with Tukey's multiple comparisons test: *RAX*: noBMPvsd5-14 ***p= 0.0007, d5-7vsd5-14 **p= 0.0069, noBMPvsd5-11 ***p= 0.0003, d5-7vsd5-11 **p= 0.0028, noBMPvsd5-9 *p= 0.0372, POMC: d5-11vsd5-14 *p=0.0166. **c**, ICC of TRH in d50 ARC RC17 3D cultures of the BMP withdrawal experiment. **d**, qRT-PCR of d16 ARC progenitors derived from a hiPSC line (KOLF2.1) with different termination days of BMP4 treatment (n=1). **e**, qRT-PCR of d50 ARC culture derived from a hiPSC line (KOLF2.1) for markers for ARC (*AGRP*), VMH (*GPR149*), and tanycytes (*RAX, FGF10)* (n=1). **f**, Combinatorial immunocytochemistry (ICC) (NKX2-1) and *in situ* hybridisation (ISH) (*RAX, TBX3*) on d16 ARC cultures derived from a hiPSC line (KOLF2.1). Scale bars: 100 µm. BMP, bone morphogenetic protein; VMH, ventromedial hypothalamus.
